# BRD4, Mediator, and Pol II form heterogeneous condensates with distinct transcriptional and acetylation-dependent states

**DOI:** 10.64898/2026.06.04.729043

**Authors:** Samuel Shoup, Alina Schaaf, Kathrin Hertäg, Amy-Sue Sattler, Márton Gelléri, Fridolin Kielisch, Thomas Speck, Sandra Schick

## Abstract

Transcriptional condensates at super-enhancers are thought to concentrate BRD4, Mediator, and RNA polymerase II (Pol II) to promote gene activation, yet their compositional organization and regulation remain poorly understood. We developed a high-throughput live-cell phenomics platform based on endogenous fluorescent tagging of BRD4, MED14 (Mediator), and POLR2A (Pol II) to systematically quantify transcriptional condensate states across >1,000 chemical perturbations. Contrary to prevailing models of largely co-occupied assemblies, we find compositionally heterogenous condensate populations. In particular, BRD4-only spots emerged as a prominent class that is depleted of Mediator and Pol II, enriched at chromatin, and resistant to transcription initiation inhibition. Mechanistically, compound screening coupled to mechanism-of-action analysis identifies histone acetylation as a dominant regulatory axis for BRD4-only spots: Bromodomain and Extra-Terminal motif (BET) and histone acetyltransferase inhibition selectively deplete BRD4-only condensates, while histone deacetylase inhibition expands them. Together, these findings support a model in which acetylation-dependent BRD4 condensates define a distinct chromatin-associated regulatory state that is separable from canonical transcriptionally engaged condensates. More broadly, our work establishes condensate composition as a quantitative phenotype and provides a scalable framework for systematically dissecting the regulation of condensates across perturbations, cell types, and disease contexts.

## Introduction

Transcription is the first and a highly regulated step in gene expression, governing cellular identity, differentiation, and responses to environmental signals^1–5^. RNA polymerase II (Pol II) transcribes DNA into messenger RNA through coordination of transcription factors (TFs), co-activators including Bromodomain-containing protein 4 (BRD4) and Mediator, at regulatory chromatin elements such as enhancers and promoters^6–8^. Dysregulation of transcriptional control contributes to developmental disorders and diseases including cancer^9–11^. Within this regulatory landscape, super-enhancers, clusters of enhancers that concentrate transcriptional machinery, disproportionately control cell identity genes and are frequently dysregulated in disease contexts^9–12^. Notably, super-enhancers at oncogenes are selectively sensitive to perturbations, making them attractive therapeutic targets in cancer^12^.

The activity of super-enhancers is dependent on the acetylation state of underlying chromatin, which in turn is controlled by the opposing actions of histone acetyltransferases (HATs) and histone deacetylases (HDACs). HATs such as p300/CBP deposit acetyl marks, particularly H3K27ac, at active enhancers, whereas HDACs remove these modifications to favor a repressive chromatin state^13,14^. This dynamic acetylation landscape is read out by bromodomain (BD)-containing co-activators such as BRD4, whose binding to acetylated histones facilitates co-activator recruitment and transcription^15^. Apart from its acetylation reader and intrinsic HAT activity important for decompaction of chromatin and promoting transcription^16^, BRD4 also full-fills BD-independent functions in transcription regulation. BRD4 forms a complex with the positive transcription elongation factor b (P-TEFb; containing the catalytic subunit cyclin dependent kinase CDK9 and a regulatory cyclin subunit), which is essential for release of promoter-proximally paused Pol II into elongation and recruitment of 3′-end RNA processing factors^17,18^. The concentration of HAT activity, acetylated chromatin, and bromodomain readers at super-enhancers thus provides a molecular basis for their transcriptional potency.

The classical enhancer-promoter looping model, wherein architectural proteins bridge distal regulatory elements to gene promoters, has been refined by observations that transcriptional regulators form dynamic nuclear spots at active loci^19,20^. Many transcriptional proteins contain intrinsically disordered regions (IDRs) and engage in multivalent interactions that can drive liquid-liquid phase separation into biomolecular condensates^21–24^. This has led to the transcriptional condensate model^25^, wherein TFs, co-activators including BRD4 and Mediator, and Pol II concentrate at super-enhancers through phase separation to coordinate transcription.

Recent studies have provided evidence for transcriptional condensates through *in vitro* reconstitution, fixed-cell imaging, and perturbation experiments. BRD4 and Mediator have been shown to form condensates at super-enhancers, and Pol II clusters at active genes^26–29^. Live-cell super-resolved imaging of Mediator and Pol II revealed both unique and larger, more stable colocalizing clusters all dependent on BRD4 chromatin association; however, only Pol II stable clusters dissolved upon inhibition of the CDK9 kinase, which is important for Pol II release into transcription elongation^26^. However, a key unresolved question is whether BRD4, Mediator, and Pol II co-condense into unified assemblies or form distinct, differentially regulated populations with specialized functions. Prevailing models based on co-immunoprecipitation^28^, chromatin profiling^10^, and imaging^26,28,29^ suggest BRD4 and Mediator strongly co-occupy super-enhancers, implying near-complete spatial overlap. Yet these conclusions rely on population-averaged or overexpression-based approaches that cannot resolve single-condensate heterogeneity in living cells. This underscores the need for quantitative, endogenous, live-cell approaches that can profile condensate composition and regulation across diverse perturbation contexts without overexpression artifacts.

Here, we address these questions using endogenously tagged human cancer cells expressing fluorescently tagged BRD4, MED14 (Mediator), and POLR2A (Pol II) to enable quantitative live-cell imaging of transcriptional condensates. We first establish that these markers form partially overlapping yet compositionally distinct spot populations, with BRD4-only spots representing a previously uncharacterized large condensate class that is distinctly regulated. Through targeted perturbations of transcription initiation (CDK7 inhibition) and elongation (CDK9 inhibition), we reveal stage-coupled dependencies wherein BRD4, MED14, and POLR2A spots exhibit differential sensitivities. We then scale this approach to a high-throughput chemical screen of >1,000 compounds with mechanism-of-action (MOA) annotation, identifying histone acetylation as a dominant regulatory axis. Validation experiments using BET, HAT, and HDAC inhibitors establish that BRD4-only spots are uniquely sensitive to acetylation state and may represent enhancer-priming condensates that precede Mediator and Pol II recruitment. Together, these findings provide a quantitative framework for dissecting transcriptional condensate organization and identify acetylation-dependent BRD4-only assemblies as a foundational step in condensate-mediated transcriptional regulation.

## Results

### Endogenous tagging and live-cell microscopy pipeline enable quantitative detection of transcriptional condensates

To visualize transcriptional condensates under near-physiological conditions and avoid artifacts associated with protein overexpression, we generated endogenously tagged human HAP1 reporter cell lines. Using CRISPR-assisted insertion tagging (CRISPaint)^30^, we introduced fluorescent tags into the endogenous loci of key transcriptional machinery components in HAP1 cells (**Figure 1A**; **Supplementary Figure S1A**). Starting from a parental BRD4::eGFP line, where BRD4 is intron-tagged with eGFP, we tagged MED14 (a central scaffold Mediator subunit^31^) and POLR2A (the catalytic subunit of RNA polymerase II) with C-terminal TagRFP, generating BRD4::eGFP/MED14::TagRFP and BRD4::eGFP/POLR2A::TagRFP double knock-in lines. Genomic PCR confirmed correct integration at both loci (**Supplementary Figure S1B**), and Western blotting showed molecular weight shifts consistent with TagRFP addition (∼27 kDa) and expression levels comparable to wild-type cells (**Supplementary Figure S1C**).

**Figure 1:**
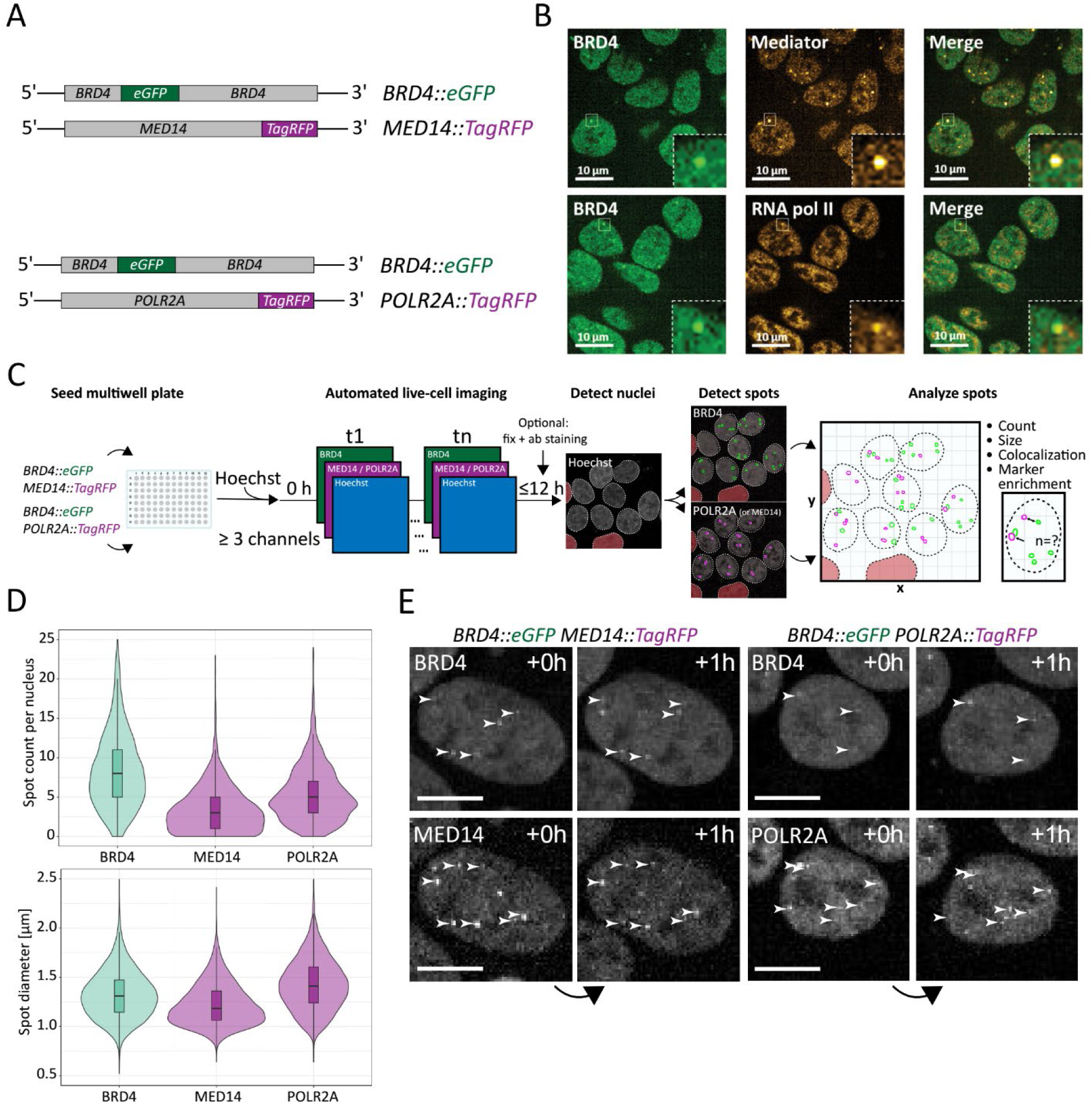
Endogenous tagging and live-cell imaging pipeline enable quantitative, longitudinal detection of transcriptional condensates. **(A)** Schematic of used endogenously tagged cell lines. TagRFP was inserted at the C-terminus of MED14 or POLR2A in the parental HAP1 BRD4::eGFP cell line, generating double-tagged BRD4::eGFP/MED14::TagRFP and BRD4::eGFP/POLR2A::TagRFP reporter lines. **(B)** Live-cell confocal images showing nuclear BRD4::eGFP signal together with MED14::TagRFP (top row) or POLR2A::TagRFP (bottom row), with merged channels. Insets show magnified examples of discrete nuclear spots. Scale bars, 10 µm. **(C)** Schematic of the automated high-throughput imaging and analysis pipeline. HAP1 reporter cells were seeded in 96- or 384-well plates, stained with Hoechst 33342, and imaged over ≤12 hours at multiple timepoints in ≥3 channels. After optional post-fixation antibody staining, an automated Harmony image analysis workflow segmented nuclei and detected spots per channel, extracting spot count, size, colocalization, and marker enrichment per nucleus. **(D)** Distributions of nuclear spot counts (top) and diameter (bottom) for BRD4, MED14, and POLR2A under 0.01% DMSO baseline conditions. Data pooled from 4 wells in 1 replicate 4 wells, which were checked to have similar distributions, were pooled. (E) Representative time-lapse confocal images (z-stack, maximum intensity projection) of BRD4::eGFP/MED14::TagRFP (left) and BRD4::eGFP/POLR2A::TagRFP (right) nuclei at t = 0h and t = +1h. Arrowheads indicate spots that remained positionally stable over the interval, demonstrating that spots are not transient aggregates. Scale bars, 5 µm.

Live-cell confocal imaging revealed that BRD4, MED14, and POLR2A each formed large, punctate nuclear spots (**Figure 1B**). These observations are consistent with previous reports demonstrating that BRD4, Mediator, and Pol II organize into dynamic nuclear spots at active regulatory regions in other cell types and organisms^26–29^. To enable systematic, quantitative analysis of condensate dynamics across diverse experimental conditions, we developed a high-throughput live-cell imaging and analysis platform (**Figure 1C**). Cells were seeded in 96- or 384-well plates, stained with Hoechst 33342 to label chromatin, and imaged on an automated confocal microscope over 12 hours. An automated image analysis workflow segmented nuclei and detected spots in each channel, extracting parameters including spot count, diameter, colocalization metrics, marker enrichment, and nearest-neighbor distances between spot markers. This approach enabled single-nucleus and single-spot quantification of condensate features across thousands of cells per condition.

Under baseline conditions using confocal microscopy, BRD4, MED14, and POLR2A displayed distinct spot count distributions per nucleus (**Figure 1D**). Interestingly, BRD4 spots were more abundant (median ∼8 spots/nucleus) than MED14 (∼3 spots/nucleus) or POLR2A (∼5 spots/nucleus), and spot sizes ranged from ∼1–1.5 μm in diameter for all three markers. These distributions are consistent with prior studies reporting varying spot numbers and sizes across individual nuclei, and contrast with studies using overexpressed IDR constructs that often detect far more spots due to artificially elevated protein concentrations.

Notably, the spots measured in confocal microscopy likely represent the upper tier of a broader size continuum. Super-resolution single-molecule localization microscopy (SMLM) revealed the presence of many additional, smaller sub-diffraction clusters not detectable by conventional confocal imaging (**Supplementary Figure S1D, E**). These sub-diffraction structures showed an exponential decay in diameter distribution starting from ∼100 nm, with a secondary, less abundant population at 300–400 nm likely corresponding to the larger spots visible in confocal images. The apparent enlargement of spots in confocal images likely reflects the combined effects of the microscope point spread function and threshold-dependent automated segmentation.

To address whether these microscopic spots represent stable structures or transient aggregates, we performed time-lapse imaging and fluorescence recovery after photobleaching (FRAP) experiments. Individual BRD4::eGFP spots exhibited rapid fluorescence recovery (t₁/₂ ≈ 2.8 ± 0.2 s, mobile fraction ≈ 81.6 ± 3.6 %; **Supplementary Figure 1F**), indicating high molecular turnover consistent with a dynamic condensate. Still, many spots remained visible and relatively positionally stable for one hour or longer (**Figure 1E**).

Together, this suggests that these are the longer-lived liquid-like condensates previously described in literature^26,27,29^. The combination of dynamic molecular exchange and prolonged structural stability justifies longitudinal time-course imaging, enabling longitudinal tracking of condensate responses to perturbations. Overall, these data establish endogenous reporter lines and a high-throughput imaging framework for quantitative condensate profiling.

### BRD4, Mediator, and Pol II form heterogeneous condensate populations, including a BRD4-only subpopulation depleted of Mediator and Pol II

Having established quantitative detection of BRD4-, MED14-, and POLR2A-defined spots, we next determined their spatial relationships and molecular composition. Previous studies suggest that the co-activators BRD4 and Mediator are highly colocalizing^12,29^, while co-activators and Pol II can form distinct condensates with varying CTD phosphorylation states^32–34^. However, quantitative profiling of their spatial overlap and functional signatures at endogenous expression levels across diverse conditions has remained limited.

We first quantified colocalization by defining overlapping spots as those with center-to-center distances less than the sum of their radii (**Figure 2A**). Surprisingly, only ∼20% of BRD4 spots colocalized with MED14 and ∼30% with POLR2A, whereas ∼50% of MED14 spots and ∼40% of POLR2A spots overlapped with BRD4 (**Figure 2B**). To test whether this exceeded random spatial association, we compared observed colocalization to randomized distributions (**Figure 2C**). All marker pairs showed higher-than-random colocalization, with BRD4↔MED14 pairs trending to slightly higher enrichment than BRD4↔POLR2A pairs, consistent with partial spatial segregation of co-activator and elongation condensates. Rather than forming a unified assembly, BRD4, MED14, and POLR2A can occupy non-overlapping spot populations, with only some BRD4 spots colocalizing with Mediator or Pol II in living cells.

**Figure 2:**
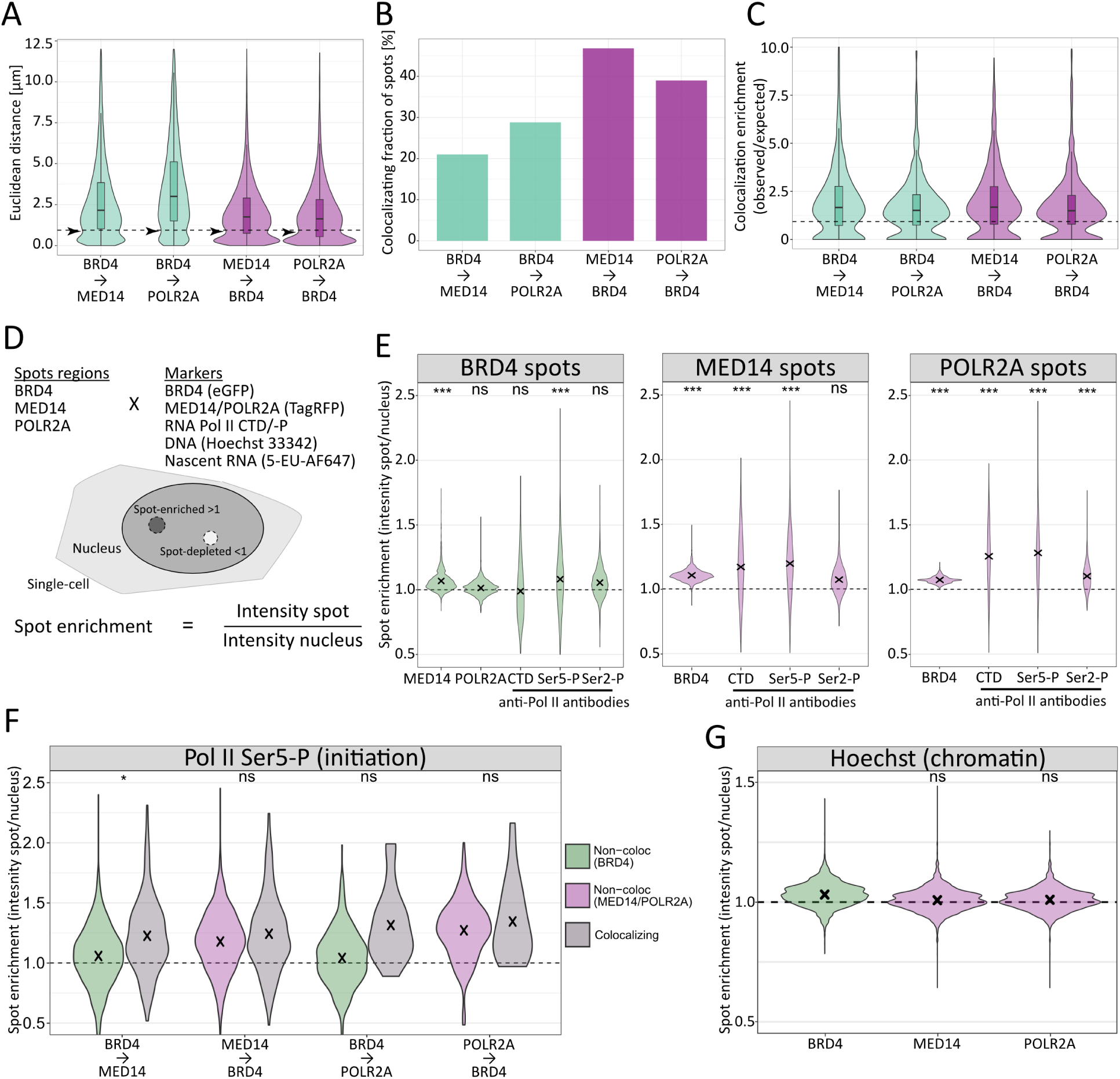
BRD4 condensates are compositionally heterogeneous: a BRD4-only subpopulation is depleted of Mediator, Pol II, and Pol II CTD phosphorylation marks. **(A)** Violin plots of pairwise minimum Euclidean distances [µm] between nearest spot centers for all four directionality pairs (BRD4→MED14, BRD4→POLR2A, MED14→BRD4, POLR2A→BRD4), computed across >100,000 spots per timepoint under DMSO baseline conditions. Dashed line indicates average spot diameter and arrows indicate that this corresponds to a dip in distance frequency. **(B)** Bar plots showing the colocalizing fraction of spots [%] for each marker pair. Two spots were defined as colocalizing when their center-to-center distance was less than the sum of their radii. **(C)** Colocalization enrichment (observed/expected ratio) comparing actual colocalization frequencies to those from a random spot placement simulation in which spot number per nucleus was preserved but positions were randomized. Dashed line at 1 indicates random expectation. A-C data were pooled from 4 separate wells of one replicate per cell line, which were similarly distributed. **(D)** Schematic of the single-cell spot enrichment metric (mean spot intensity / mean nuclear intensity). Values >1 indicate spot enrichment; values <1 indicate depletion. Markers quantified include BRD4::eGFP, MED14::TagRFP or POLR2A::TagRFP, RNA Pol II CTD and phosphorylated forms (immunofluorescence after fixation), chromatin (Hoechst 33342). **(E)** Spot enrichment within BRD4 spots (left), MED14 spots (center), and POLR2A spots (right). Markers are indicated at the bottom. Dashed line at 1, indicating no enrichment relative to the nucleoplasm. Violins show the single-cell distribution; cross (×) indicates the mean. Data were aggregated to the replicate level prior to statistical testing. One-sample t-tests against µ = 1 with Holm correction for multiple comparisons within each spot type. **(F)** Ser5-P enrichment (spot intensity / nucleus intensity) within BRD4 spots stratified by colocalization status (non-colocalizing BRD4-only spots (green), MED14 or POLR2A spots non-colocalizing with BRD4 (purple) vs. colocalizing BRD4+MED14 or BRD4+POLR2A spots (grey)). Violins show the single-cell distribution; cross (×) indicates the mean. Data were aggregated to the replicate level prior to statistical testing. Paired t-tests between colocalization groups with Holm correction for multiple comparisons. **(G)** Hoechst 33342 enrichment within BRD4, POLR2A, and MED14 spots. Paired t-test with Holm correction for multiple comparisons. *p < 0.05, **p < 0.01, ***p < 0.001, ns = not significant. E-G data from 2 replicates.

To determine whether the spatial heterogeneity observed above reflects distinct molecular states, we quantified enrichment of transcriptional markers within spots relative to the surrounding nucleoplasm (enrichment = spot intensity / nuclear intensity; **Figure 2D**). We focused on the C-terminal domain (CTD) of RNA Pol II, a repetitive heptapeptide sequence^35^ whose phosphorylation status serves as a hallmark of transcriptional progression^36,37^: Serine 5 phosphorylation (Ser5-P) marks initiation, while Serine 2 phosphorylation (Ser2-P) marks productive elongation. Immunofluorescence for total Pol II CTD, Ser5-P, and Ser2-P revealed distinct molecular profiles across spot populations (**Figure 2E**): BRD4 spots showed modest MED14 and Ser5-P enrichment but minimal Ser2-P, whereas MED14 spots were enriched for BRD4, total CTD, and Ser5-P but showed no significant Ser2-P enrichment, and POLR2A spots were enriched for all markers, with POLR2A spots trending toward higher CTD phosphorylation. Importantly, substantial spot-to-spot heterogeneity existed within each marker population. Together, these molecular profiles indicate that BRD4, MED14, and POLR2A spots represent compositionally heterogeneous condensate populations rather than a single, uniform transcriptional condensate type. Only a subfraction of BRD4 spots were enriched for MED14, POLR2A, and Pol II phosphorylation marks — the pattern predicted by prevailing models of BRD4-Mediator co-condensation at transcriptionally active super-enhancers — while the majority remained depleted of these transcriptional components. These BRD4 spots depleted of Mediator and Pol II activity markers are referred to from now on as BRD4-only spots. They may represent a previously under-appreciated condensate class that reflect chromatin-associated assemblies existing independently of active transcriptional machinery.

Given the incomplete colocalization between the markers and the enrichment of BRD4 spots for Ser5-P, we next asked whether the transcription initiation state of individual spots correlates with the spatial proximity of BRD4 spots to Mediator or Pol II. We therefore stratified spots based on colocalization status and compared Ser5-P enrichment in BRD4 colocalizing versus non-colocalizing populations (**Figure 2F**). BRD4 spots that colocalized with MED14 showed significantly higher Ser5-P compared to BRD4-only spots (non-colocalizing), while those colocalizing with POLR2A spots showed a trend toward higher enrichment. Interestingly, BRD4 spots also differed in their chromatin content, showing higher Hoechst signal enrichment than MED14 or POLR2A spots, which were similar to one another (**Figure 2G**). Together, these data establish that BRD4-only spots represent a distinct condensate type: depleted of Mediator, Pol II activity markers, and more enriched in chromatin.

### BRD4, MED14, and POLR2A spots respond differently to transcription stage inhibitors

The distinct CTD phosphorylation patterns and the partial spatial segregation of BRD4, MED14, and POLR2A spots suggest that these compositionally distinct structures are associated with different stages of the transcription cycle. Consistent with this notion, previous studies have proposed that Pol II transitions from transcription initiation condensates into condensates associated with transcription elongation and RNA processing^32,38^. We therefore hypothesize that the spot populations identified here represent distinct transcriptional states and would respond differentially to perturbations targeting the corresponding stage of transcription. To test this, we perturbed transcription initiation with THZ1, a CDK7 inhibitor that blocks Ser5 phosphorylation^39,40^, and elongation with Flavopiridol, a CDK9 inhibitor that prevents Ser2 phosphorylation and pause release^41,42^ (**Figure 3A**). Both compounds reduced nascent RNA synthesis measured by 5-ethynyluridine (5-EU) incorporation, with THZ1, the upstream initiation inhibitor, producing stronger suppression (**Figure 3B**), indicating effective transcriptional inhibition.

**Figure 3:**
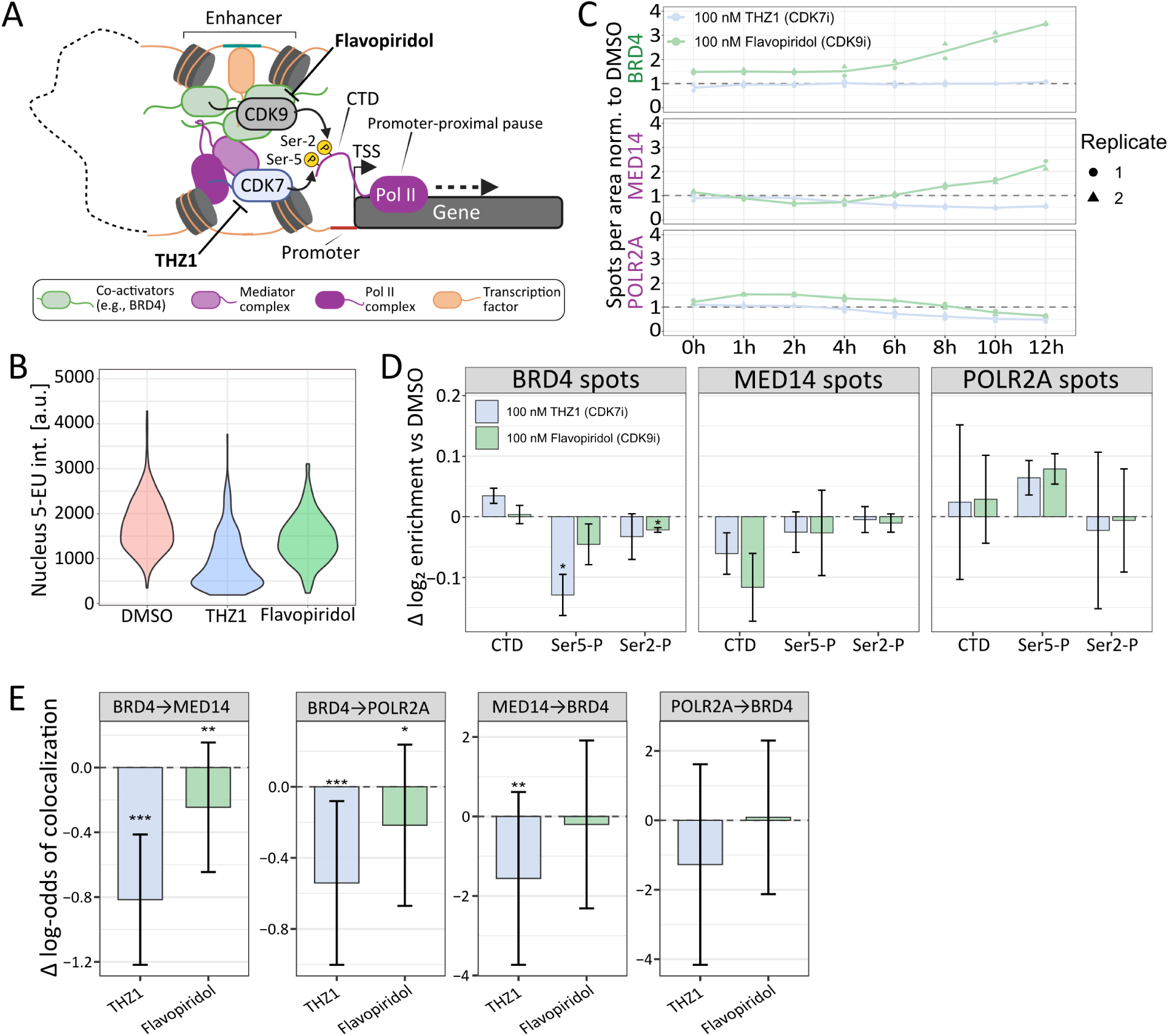
Stage-specific transcriptional inhibition reveals differential and divergent dependencies of BRD4, MED14, and POLR2A condensates. **(A)** Schematic of the transcription cycle showing CDK7-dependent Pol II CTD Ser5 phosphorylation (initiation) and CDK9-dependent Ser2 phosphorylation (pause release and elongation), with BRD4, Mediator, and Pol II depicted at enhancer and promoter. CDK7 is inhibited by THZ1 (100 nM); CDK9 by Flavopiridol (100 nM). Figure was created using BioRender and image processing software. Created in BioRender. Schick, S. (2026) https://BioRender.com/n62pu2a **(B)** Violin plots of nucleus 5-EU fluorescence intensity (arbitrary units, a.u.) after 12 hours of treatment with 0.01% DMSO, 100 nM THZ1, or 100 nM Flavopiridol. Data is pooled from multiple wells from 1 replicate. **(C)** Spot number per nuclear area for BRD4, MED14, and POLR2A over 0–12 hours under 0.01 % DMSO, 100 nM THZ1, and 100 nM Flavopiridol treatment, normalized to the DMSO median at each timepoint. Data points indicate replicate means (n = 2 independent replicates). **(D)** Bar charts showing log₂(enrichment_treatment / enrichment_DMSO) for the indicated markers (CTD, Ser5-P, Ser2-P) within BRD4 spots (left), MED14 spots (center), and POLR2A spots (right) at the 12-hour endpoint. Enrichment is defined as mean intensity within spots divided by mean nuclear intensity. Δ log₂ values were computed at the replicate level; bars show the mean, error bars ± 1 standard deviation (SD) across replicates. Statistical comparisons by one-sample t-test of Δ log₂ vs. 0; p-values Holm-corrected within each spot type × antibody panel. **(E)** Bar charts showing Δ log-odds ratio of colocalization treatment versus DMSO for all four marker pair directionalities (BRD4→MED14, BRD4→POLR2A, MED14→BRD4, POLR2A→BRD4) under 100 nM THZ1 and 100 nM Flavopiridol treatment. The log-odds ratio was computed per field as logit(p_coloc_treatment) − logit(p_coloc_DMSO), with logit(p) = log(p / (1 − p)) and p clipped to [0.001, 0.999] to avoid infinite values. Only fields with ≥ 5 spots were included. Bars show the mean Δ log-odds, error bars ± 1 SD across fields. Statistical comparisons by one-sample t-test of Δ log-odds vs. 0; p-values Holm-corrected within each direction. For all panels, n = 2 replicates. *p < 0.05, **p < 0.01, ***p < 0.001, ns = not significant.

To assess the stage-specific dependence of condensate populations, we quantified spot abundance over 12 hours after inhibitor treatment (**Figure 3C**). BRD4 spots persisted under the CDK7 inhibitor THZ1, maintaining counts comparable to DMSO-control baseline throughout the timecourse. This persistence indicates that a substantial fraction of BRD4 spots can be maintained independently of active transcription initiation. In contrast, MED14 spots and POLR2A spots declined to approximately half of baseline after 6 hours of THZ1 treatment and stayed reduced, consistent with a strong reliance on ongoing initiation for their stability. Treatment with the CDK9 inhibitor Flavopiridol produced a strikingly different pattern: POLR2A spots initially increased within the first hours – reflecting likely an accumulation of promoter-proximal, paused Pol II - before declining at later timepoints, whereas both BRD4 and MED14 spots accumulated as POLR2A spot numbers decreased. These divergent kinetics demonstrate that spot maintenance is not uniformly tied to transcription but instead reflects factor-specific coupling to distinct transcriptional stages: BRD4 can persist without transcription initiation, whereas Mediator and Pol II are primarily initiation-dependent, and their balance shifts markedly when elongation is blocked.

To determine whether stage-specific inhibitors alter the molecular composition of remaining spots rather than only their abundance, we quantified changes in Pol II CTD, Ser5-P, Ser2-P within BRD4, MED14, and POLR2A spots at the 12-hour endpoint (**Figure 3D**). While MED14 and POLR2A spots did not show any significant effects, BRD4 spots were selectively reduced in Ser5-P enrichment after THZ1, consistent with loss of initiation activity. Ser2-P remained near baseline and showed a small reduction after Flavopiridol. Spatial relationships between markers were similarly disrupted (**Figure 3E**): THZ1 reduced colocalization between BRD4 and both MED14 and POLR2A spots. These data indicate that inhibition of transcription initiation reduces BRD4 spots colocalizing with Mediator/Pol II, while increasing BRD4-only spots.

Together, these data establish that BRD4, MED14, and POLR2A spots exhibit stage-coupled dependencies. Upon initiation inhibition, half of the MED14 and POLR2A spots collapse while BRD4 spots are maintained. This goes along with an increase in number of BRD4-only spots and a decrease in colocalizing spots. After elongation inhibition, BRD4 and MED14 spots are increased, while a subpopulation of POLR2A spots is lost. POLR2A stalls at the respectively modified stage, which may go along with their retention in transcription initiation condensates. These data support a model wherein these transcriptional regulators form different condensates that are functionally specialized for distinct transcriptional stages.

### A live-cell imaging screen quantifies BRD4/POLR2A condensate phenotypes across >1,000 compounds

To systematically identify which cellular processes beyond CDK7 and CDK9 modulate condensate organization and affect the different condensate types, we extended our high-throughput imaging pipeline to a large-scale compound screen. We screened more than 1,000 compounds from four different libraries: epigenetic modulators, kinase inhibitors, anti-cancer agents, and the CeMM Library of Unique Drugs (CLOUD)^43^, arrayed across ten 384-well plates in two concentrations per compound (**Figure 4A**). Each plate included matched controls: DMSO vehicle as negative control, and three different positive control compounds. Due to technical constraints, the compound screen was conducted in a single replicate; consequently, the data are presented as an initial screening dataset.

**Figure 4:**
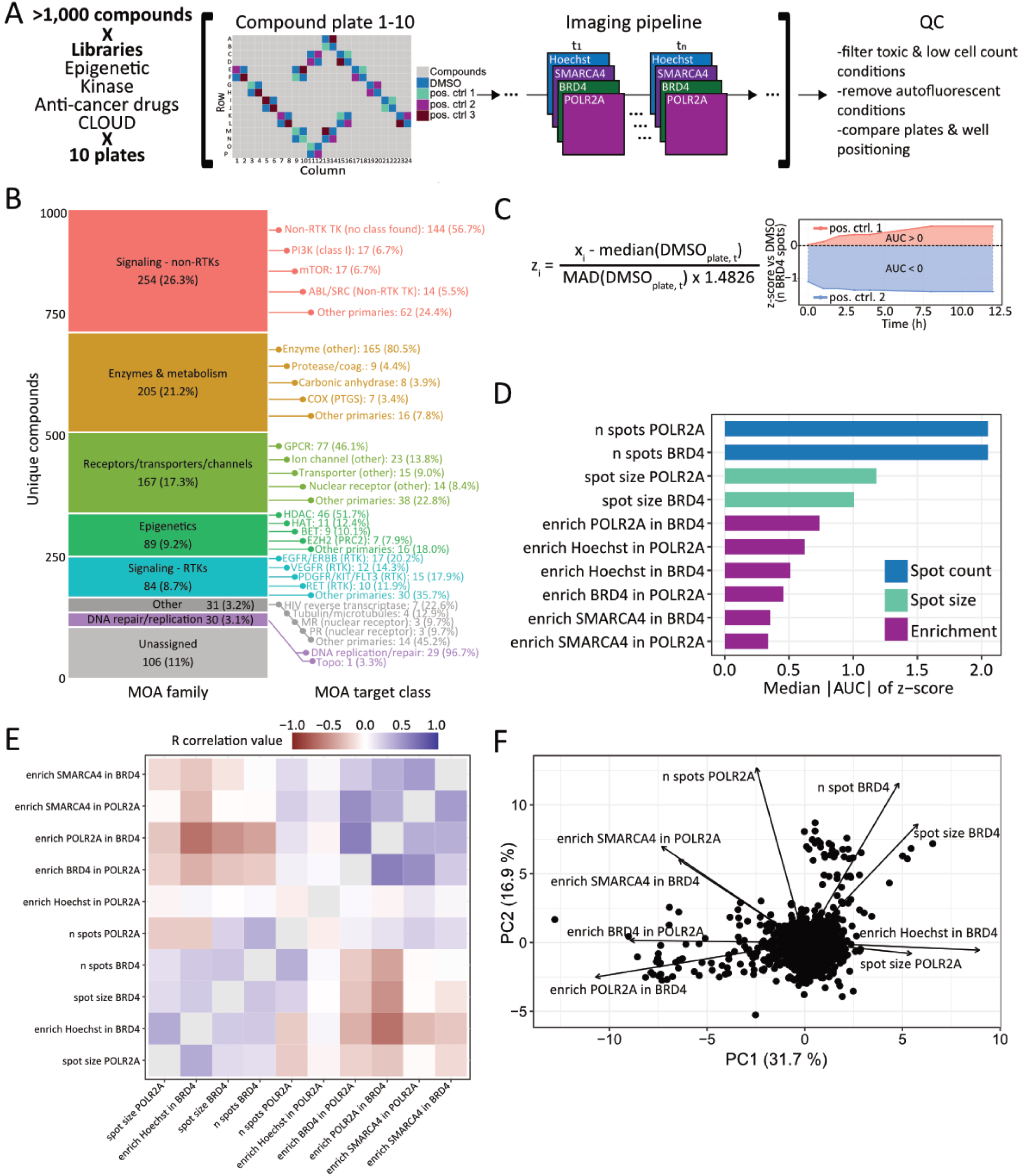
High-content live-cell screening of >1,000 compounds reveals structured, time-resolved condensate phenotypes. **(A)** Compound screen design schematic. BRD4::eGFP/POLR2A::TagRFP/SMARCA4::SNAP-dTAG-HA HAP1 cells were arrayed across ten 384-well plates, testing >1,000 compounds from four libraries (Epigenetic, Kinase, Anti-cancer, CLOUD) at two concentrations per compound. Each plate included DMSO as vehicle control and three positive controls. Cells were imaged over 12 hours at seven timepoints (0, 1, 2, 3, 4, 8, 12 h), acquiring two fields per well (∼100 nuclei per well). A QC workflow removed wells with low cell counts, toxicity, or autofluorescence artifacts; plates 9 and 10 were excluded due to globally elevated nuclear intensities. **(B)** Distribution of screened compounds by mechanism of action (MOA) family (stacked bar) and MOA target class (outer labels), derived from PubChem- and ChEMBL-based annotation. 11% of compounds remained unassigned. **(C)** Robust Z-score normalization applied per plate per timepoint. Representative Z-score time courses for BRD4 spot counts under positive controls illustrate the signed area-under-the-curve (AUC) summarization. The scaling factor 1.4826 is derived from the reciprocal of the 75^th^ percentile of the standard normal distribution. AUC values could further be expressed as AUC% by scaling to the per-feature dynamic range (max AUC = 100%). **(D)** Ranked median absolute AUC of z-score across the ten extracted features, colored by feature category (spot count (blue), spot size (green), enrichment (purple)). **(E)** Unsupervised hierarchically clustered pairwise Pearson correlation heatmap of the ten features across all compound-concentration conditions. **(F)** PCA loadings plot of the ten-feature AUC matrix across all 2,154 compound-concentration conditions. Arrows indicate the direction and magnitude with which each feature drives variance in PC space. PC1 (31.7% variance explained) and PC2 (16.9% variance explained).

For the screen, we expanded our dual-tagged cell line to include SMARCA4::SNAP-dTAG-HA (BRG1, the main catalytic ATPase subunit of BAF/SWI-SNF chromatin remodeling complexes in HAP1 cells). BAF complexes are recruited to active regulatory regions where it regulates chromatin accessibility required for transcription factor binding and Pol II activity^44–47^. Recent studies have demonstrated that BAF complex subunits contain IDRs that drive condensate formation and recruitment to transcriptional hubs, with oncogenic fusions and pioneer factors exploiting these properties to redirect chromatin remodeling activity^48–50^. Unlike BRD4, MED14, and POLR2A, SMARCA4 did not form discrete microscopic spots in our system but showed diffuse nuclear localization with variable enrichment at BRD4 and POLR2A spots. We therefore quantified SMARCA4 as an enrichment metric within BRD4/POLR2A spots rather than spot counts, serving as a readout of chromatin remodeler recruitment to condensates and enabling assessment of how perturbations may affect chromatin remodeling activity within spots.

For the high-throughput compound screen, we imaged BRD4::eGFP/POLR2A::TagRFP/SMARCA4::SNAP-dTAG-HA reporter cells over 12 hours at seven timepoints, extracting spot counts, sizes, marker enrichments, and spatial relationships through the automated analysis pipeline established previously. To ensure data quality across this multi-plate experiment, we implemented stringent quality control filters. Plates 9 and 10 showed globally elevated nuclear intensities and were removed entirely (**Supplementary Figure S2A**). Wells with fewer than 10 starting nuclei, evidence of toxicity (low nuclei ratio at 12h/0h), or compound autofluorescence (top/bottom intensity outliers) were excluded (**Supplementary Figure S2B**). Across the eight retained plates containing 2,888 wells, 8.6% (n=247) failed at least one criterion, with toxicity accounting for the majority (n=153; **Supplementary Figure S2B-C**). The remaining 2641 high-quality conditions showed no systematic well-positional bias (**Supplementary Figure S2D**) and comparable DMSO baseline spot distributions across plates (**Supplementary Figure S2E**), confirming robust plate-to-plate consistency.

To enable systematic analysis by mechanism of action (MOA), we next annotated compound targets using a custom bioinformatics pipeline (**Figure 4B**). As most screened compounds lacked standardized MOA annotations, we extracted them from ChEMBL^51^ mechanisms or bioactivity data. We collapsed individual targets into MOA target classes (e.g., BET: bromodomain and extra-terminal domain proteins; HDAC: histone deacetylases; EGFR/ERBB: epidermal growth factor receptor family; PI3K: phosphoinositide 3-kinase) and further grouped these into higher-level MOA families (e.g., epigenetics, receptor tyrosine kinases, non-receptor tyrosine kinase (non-RTK) signaling, enzymes & metabolism). This unified annotation covered nearly all screened compounds, with 11% of compounds remaining unassigned. The annotated dataset revealed diverse coverage across MOA families, enabling MOA-level enrichment and signature analyses.

To enable comparison across plates, markers, and timepoints, we normalized all features by z-scoring relative to the plate-matched DMSO median and median absolute deviation (MAD) at each timepoint (**Figure 4C**). To condense 12-hour trajectories into a single comparable metric emphasizing sustained effects over transient fluctuations, we calculated the signed area under the curve (AUC) for each feature-condition pair (positive controls exemplified in Figure 4C). Positive AUC values indicate sustained increases relative to DMSO, while negative values reflect sustained decreases.

We focused analysis on ten primary features: spot counts and sizes for BRD4 and POLR2A, co-enrichment of POLR2A in BRD4 spots and vice versa, chromatin (Hoechst) enrichment in BRD4 and POLR2A spots, and SMARCA4 enrichment in both spot types as a readout of chromatin remodeler recruitment. Ranked by median absolute AUC across all conditions, spot counts for BRD4 and POLR2A exhibited the strongest responses, followed by spot sizes, while marker enrichment features showed more modest effects after the perturbations (**Figure 4D**). This hierarchy indicates that the abundance of condensates is the most dynamically regulated feature, whereas changes in geometry and molecular composition are subtler. Consequently, spot formation and dissolution appear to constitute the primary regulatory layer, while enrichment alterations may reflect secondary, possibly fine-tuning compositional remodeling.

To assess whether distinct features capture independent or redundant information, we examined pairwise correlations across all AUC values (**Figure 4E**). Hierarchical clustering revealed two major anti-correlated modules: spot count and size features clustered together and positively correlated with each other for BRD4 and negatively for Pol II. On the contrary, marker enrichment features formed a second cluster that inversely correlated with BRD4 spot abundance. This inverse relationship suggests a trade-off wherein conditions promoting many or large spots dilute marker co-enrichment, while conditions reducing spot numbers concentrate remaining markers. Notably, BRD4 and POLR2A spot counts were highly correlated, indicating that most perturbations affect both markers similarly, whereas enrichment features showed marker-specific patterns. For example, BRD4 spot counts were positively correlated with Hoechst enrichment in BRD4 spots, but this was not the case for POLR2A spots and Hoechst enrichment in POLR2A spots.

To test whether compounds separate by their phenotypic responses, we performed principal component analysis (PCA) on the ten-feature AUC matrix (**Figure 4F**). Principal component 1 (PC1) (31.7% variance) primarily captured marker enrichment features, while PC2 (16.9%) was driven by spot abundance and size. Some compounds distributed broadly across PC space further away from the cluster of most compounds in the center, indicating diverse phenotypic responses.

Together, these quality control, MOA annotation, and normalization procedures established a robust, time-resolved framework for quantifying transcriptional condensate phenotypes across thousands of chemical perturbations. The structured relationships among features, particularly the spot count/enrichment trade-off and feature-driven clustering in phenotypic space, provide a foundation for identifying convergent regulators of condensate organization in subsequent analyses. This platform demonstrates several methodological advances: endogenous tagging eliminates overexpression artifacts, live-cell imaging captures temporal dynamics, and MOA-based clustering identifies convergent regulators. The approach is scalable to other condensate systems, cell types, and perturbation modalities including genetic screens and optogenetic tools.

### Mechanism-of-action enrichment and phenotypic signatures reveal convergent regulators of condensates

Having established normalized, time-resolved condensate phenotypes across >1,000 compounds, we next sought to identify which cellular pathways most strongly regulate condensate organization. To systematically define perturbations, we ranked all conditions by their AUC values for each of the ten features and classified the top and bottom 1% as hits (**Figure 5A**, **Supplementary Figure S3**). This yielded 242 hit conditions from 2,154 screened wells, with 92 hits occurring only in the top 1%, 93 only in the bottom 1%, and 57 appearing in both top and bottom 1% across different features (**Figure 5B**). The latter group indicates compounds that drove distinct features in opposite directions—for example, reducing spot counts while increasing marker enrichment—consistent with the anti-correlated feature modules identified previously (Figure 4E-F).

**Figure 5:**
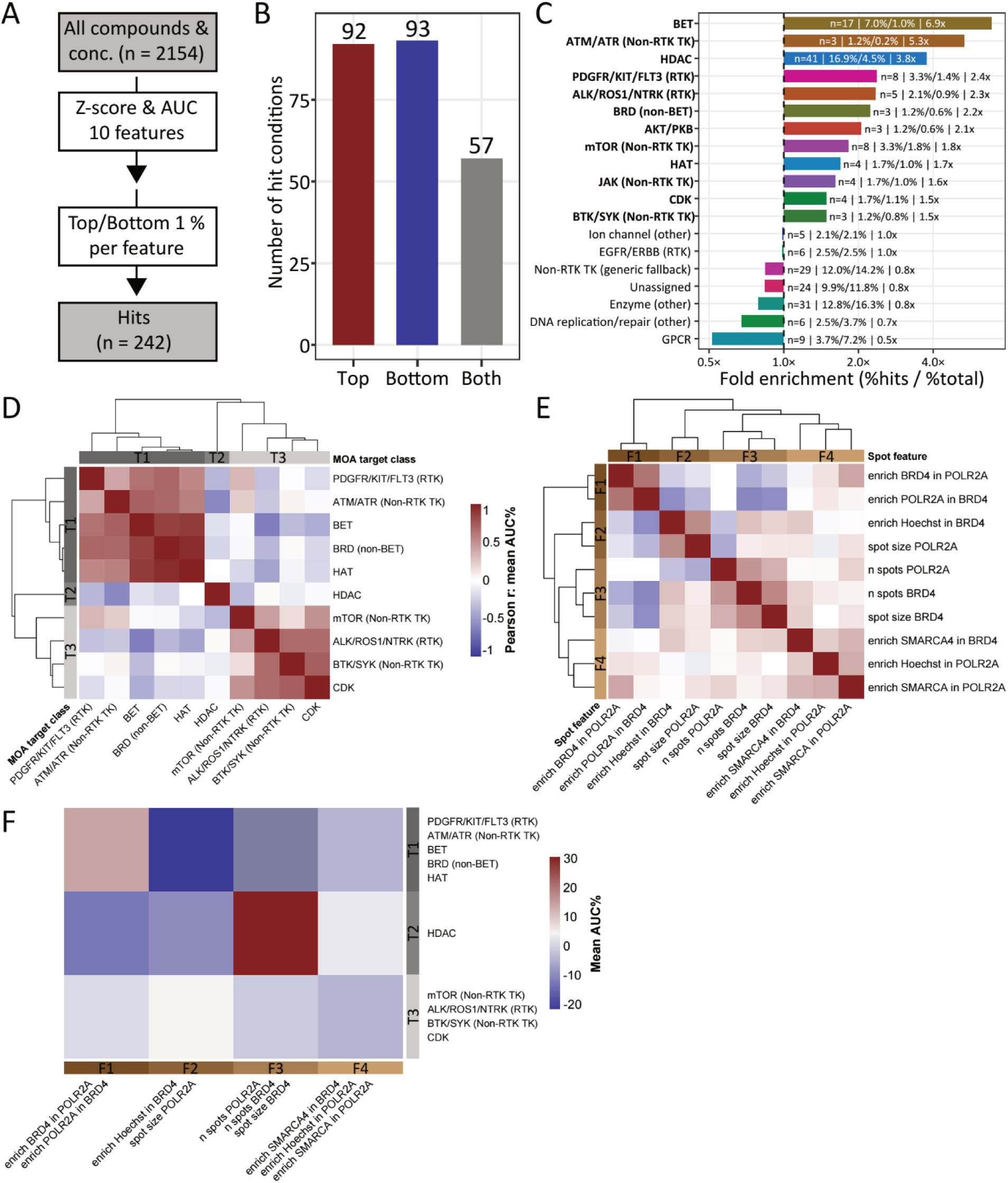
MOA enrichment and phenotypic clustering identify convergent regulators of transcriptional condensate organization. **(A)** Hit-calling workflow. All 2,154 compound-concentration conditions were ranked by signed AUC% for each of the ten features; conditions in the top or bottom 1% per feature were designated hits, yielding 242 hit conditions. **(B)** Bar chart showing the number of hit conditions falling in the top 1% only, bottom 1% only, or both top and bottom 1% across different features. **(C)** Fold enrichment of MOA target classes among hit conditions (% of MOA among compound hits / % of that MOA among total compounds screened), restricted to classes with ≥3 hit conditions. Dashed line at 1 indicates no enrichment. n=number of compounds from that MOA class amongst the total 242 hits | % amongst hits / % amongst total compounds | fold change (% amongst hits / % amongst total). **(D)** Unsupervised hierarchical clustering of MOA target classes based on Pearson correlations of their mean AUC% feature signatures, colored by Pearson r. Three MOA clusters are indicated (T1–T3). **(E)** Unsupervised hierarchical clustering of the ten spot features based on pairwise Pearson correlations of their AUC values across all conditions, revealing four feature modules (F1–F4). **(F)** Integrated heatmap of MOA target class clusters (rows, T1–T3) versus feature modules (columns, F1–F4), colored by mean AUC%.

To account for the fact that not all compound classes have the same number of represented members in the compound libraries, we calculated MOA enrichment by dividing each target class’s proportion among hits by its proportion in the total screened set (**Figure 5C**). BET inhibitors ranked first with nearly 7-fold enrichment, followed by ATM/ATR kinase inhibitors (5.3-fold) and HDAC inhibitors (3.8-fold). Additional enriched classes included HAT inhibitors, non-BET bromodomain (BRD) inhibitors, and several RTK and non-RTK kinase families including for example PDGFR/KIT/FLT3, ALK/ROS1/NTRK, AKT/PKB, and CDK. In contrast, broader or poorly defined MOA classes such as GPCR modulators and unassigned compounds were under-represented, indicating that strong condensate phenotypes were largely confined to mechanistically focused inhibitor classes. This enrichment pattern immediately highlighted epigenetic regulators, particularly those controlling histone acetylation, and kinases as dominant modulators of transcriptional condensate organization. This underscores that the screen worked and can identify regulators and their impact on the spots.

To identify groups of functionally related perturbations that elicit comparable spot phenotypes, we performed unsupervised hierarchical clustering of MOA target classes based on correlations of their feature signatures (**Figure 5D**). We classified three principal MOA clusters, termed Target 1-3 (T1-3). T1 comprised six classes—BET, HAT, non-BET BRD, ATM/ATR, PDGFR/KIT/FLT3, and EGFR/ERBB inhibitors, with BET, HAT, and BRD inhibitors forming a tightly correlated subgroup. T2 consisted solely of HDAC inhibitors, which showed no positive correlation with any other MOA class. T3 encompassed four kinase-associated classes: mTOR, ALK/ROS1/NTRK, BTK/SYK, and CDK inhibitors. Importantly, this MOA-level clustering recapitulated patterns observed in compound-level hierarchical clustering (**Supplementary Figure S4A**), confirming that shared mechanisms produce convergent phenotypes.

To systematically relate MOA responses to condensate features, we first performed hierarchical clustering on the ten features themselves (**Figure 5E**). Here we classified four feature (F) modules: F1 captured BRD4-POLR2A cross-enrichment features, F2 grouped spot size of POLR2A and chromatin (Hoechst) enrichment in BRD4 spots, F3 contained spot count features and BRD4 spot size, and F4 comprised SMARCA4 enrichment and chromatin (Hoechst) content in POLR2A spots. Integrating these clusters into a unified heatmap linkes MOA target class clusters to feature modules (**Figure 5F**, **Supplementary Figure S4B**). This analysis revealed epigenetic regulators as the strongest condensate modulators. BET, HAT, and non-BET bromodomain inhibitors (acetyl-reader/writer cluster T1) shared a common response: they reduced spot numbers and sizes while simultaneously increasing how much BRD4 and POLR2A co-enriched within remaining spots. HDAC inhibitors (acetyl-eraser cluster T2) produced the opposite effect—they increased overall spot numbers and sizes of BRD4 spots but reduced marker enrichment. Kinase inhibitors (kinase-associated cluster T3) showed milder effects, primarily influencing SMARCA4 and chromatin content in POLR2A spots.

Together, these patterns reveal that histone acetylation represents a dominant regulatory axis for condensate organization. This antagonism between acetyl-writer (HAT)/ acetyl-readers (BET/BRDs) and acetyl-erasers (HDACs) reflects known acetylation biology: Blocking the writers or readers mimics loss of acetylation, while blocking the erasers increases acetylation. Kinase perturbations, by contrast, produced weaker effects. These findings establish histone acetylation as a key regulator of transcriptional condensate organization, however, the screen-wide analysis aggregated responses across all conditions, leaving unclear whether acetylation specifically controls distinct condensate populations.

### BET/HAT/HDAC perturbations uncover acetylation-sensitive BRD4-only condensates

The antagonistic responses of BET/HAT versus HDAC inhibitors suggest that acetylation state directly controls BRD4 condensate formation. To dissect this regulation at higher resolution and test whether the acetylation axis specifically governs BRD4-only or with MED14- or POLR2A-colocalizing BRD4 spots, we performed targeted validation experiments using representative compounds from each node of the reader–writer–eraser triad: JQ1 (BET bromodomain inhibitor)^52^, A485 (p300/CBP HAT catalytic inhibitor)^53^, and SAHA/Vorinostat (pan-HDAC inhibitor)^54^ (**Figure 6A**). We imaged BRD4::eGFP/MED14::TagRFP and BRD4::eGFP/POLR2A::TagRFP cells over 12 hours to simultaneously track BRD4 spots and assess their spatial relationships with Mediator and Pol II.

**Figure 6:**
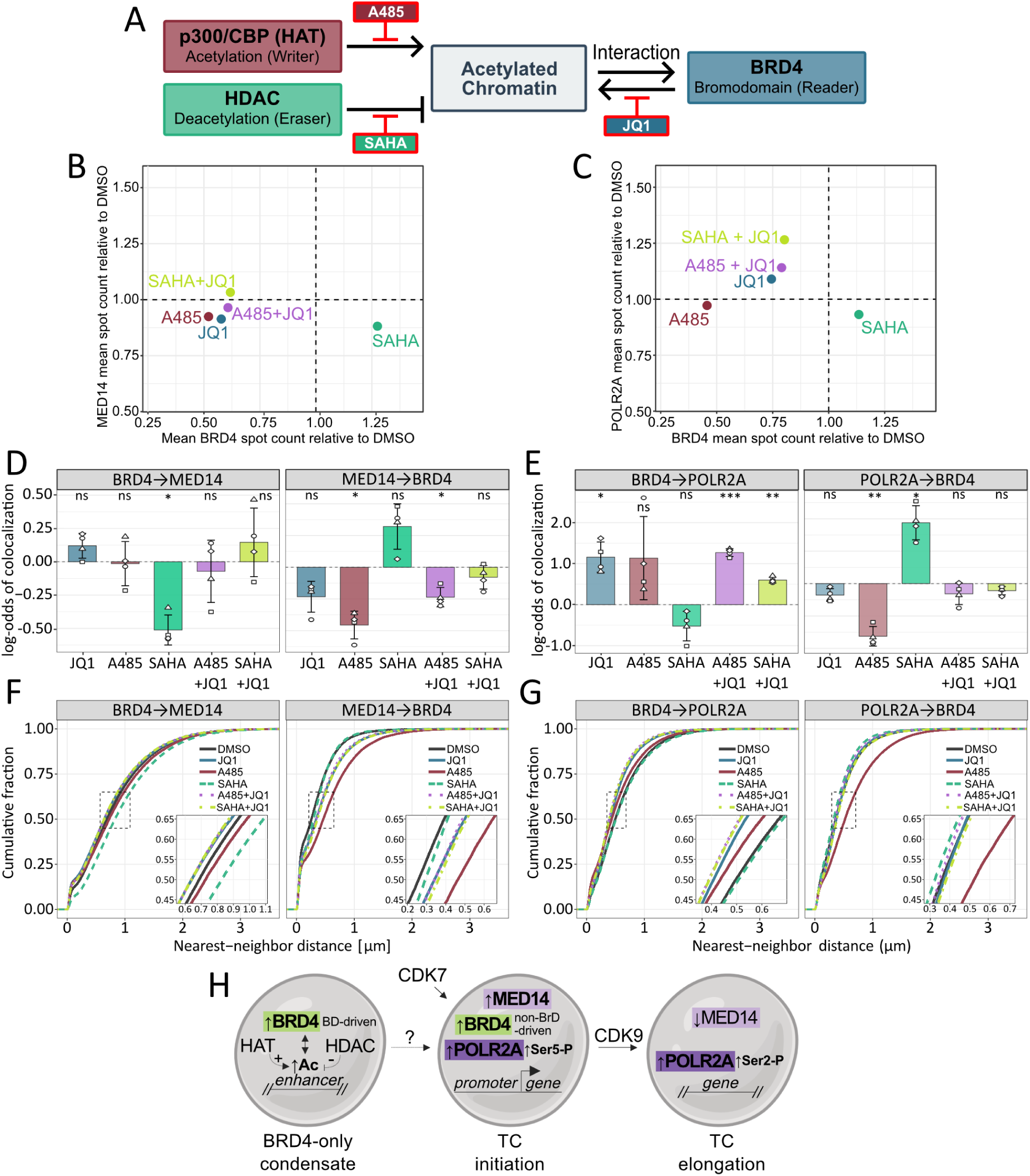
The histone acetylation reader-writer-eraser axis primarily controls BRD4-only condensates. **(A)** Schematic of the chromatin acetylation axis illustrating the interplay between the HAT writer p300/CBP, BRD4 bromodomain reader, and HDAC eraser, and their respective inhibitors. **(B)** Scatter plot of mean MED14 spot count relative to DMSO (y-axis) versus mean BRD4 spot count relative to DMSO (x-axis) in BRD4::eGFP/MED14::TagRFP cells. Each point represents the mean across fields for one treatment condition at one timepoint (0–12 h); dashed lines at 1 indicate DMSO baseline. Treatments: JQ1 (500 nM), A485 (500 nM), SAHA (500 nM), A485+JQ1 (500 nM each), SAHA+JQ1 (500 nM each). n=4 replicates pooled as mean. **(C)** As in (B), for mean POLR2A spot count relative to DMSO (y-axis) versus BRD4 spot count relative to DMSO (x-axis) in BRD4::eGFP/POLR2A::TagRFP cells. **(D)** Bar charts showing Δcolocalization (log-odds ratio of colocalization fraction relative to DMSO, logit(treatment) − logit(DMSO); mean ± SD, n=4 replicates indicated by different shaped points) for BRD4→MED14 (left) and MED14→BRD4 (right). Statistical comparisons by two-sided one-sample t-test of Δcolocalization vs. 0; p-values Holm-corrected across treatments within each direction. *p < 0.05, **p < 0.01, ***p < 0.001, ns = not significant. Different point shapes indicate replicates. **(E)** As in (D), for BRD4→POLR2A (left) and POLR2A→BRD4 (right). **(F)** Cumulative fraction plots of nearest-neighbor distance [µm] for BRD4→MED14 (left) and MED14→BRD4 (right) under all indicated conditions. Insets show zoomed distributions at the divergence region. n=4 replicates. **(G)** As in (F), for BRD4→POLR2A (left) and POLR2A→ BRD4 (right). **(H)** The data suggests a model that BRD4-only condensates constitute an acetyl-sensitive condensate class distinct from transcription initiation and elongation condensates. Figure was created using BioRender and image processing software. Created in BioRender. Schick, S. (2026) https://BioRender.com/vwe11cj

As predicted from the screen, JQ1 and A485 produced similar effects on spot abundance: both reduced BRD4 spot counts to approximately 50% of DMSO baseline by 12 hours, while SAHA increased BRD4 spots by ∼25% (**Figure 6B**). To investigate whether the effects of tuning acetylation writer and eraser activity are dependent on the acetylation reader function of BRD4, we additionally performed co-treatments with the bromodomain inhibitor JQ1. Importantly, co-treatment with JQ1 blocked the SAHA-induced increase, indicating that bromodomain function is required for HDAC inhibition to expand the BRD4 spot population. MED14 spot counts remained largely unchanged across all treatments, whereas POLR2A spot counts showed a modest increase under JQ1 (**Figure 6C**). These divergent responses indicate that acetylation perturbations regulate BRD4 condensates while leaving Mediator and Pol II assemblies largely but not entirely unaffected, consistent with BRD4’s direct role as an acetyl-lysine reader at enhancers.

As we observed a large proportion of BRD4 spots that are non-overlapping with Mediator and Pol II (Figure 2), we next asked, if specifically these BRD4-only spots are dependent on histone acetylation. For this, we examined whether the acetylation perturbations would alter spatial relationships between BRD4 and the other markers and quantified changes in colocalization following treatment (**Figure 6D-E**). Interestingly, the effects varied depending on marker directionality. For BRD4→MED14 colocalization, SAHA decreased overlap, while JQ1 and A485 showed minimal change (**Figure 6D**). In contrast, MED14→BRD4 colocalization was reduced by both A485 and combination of A485+JQ1. These data indicate that upon increased histone acetylation, BRD4 colocalizes less with Mediator, while reduced histone acetylation or binding of BRD4 to acetylated chromatin diminishes the fraction of Mediator that remains associated with BRD4. This suggests that the association of Mediator with BRD4 depends on BRD4’s chromatin anchoring. Notably, BRD4→POLR2A colocalization increased under all treatments except SAHA, and these increases persisted in combination treatments (**Figure 6E**), indicating that non-chromatin bound BRD4 prefers to recruit/associate with Pol II spots. POLR2A→BRD4 colocalization showed an increase after SAHA treatment, but a decrease after A485 treatment. Thus, when BRD4 is less chromatin bound (A485, JQ1), it becomes more frequently colocalized with POLR2A, whereas more histone acetylation results in decreased Pol II colocalization with BRD4, but more of the Pol II spots show a colocalization with BRD4. Nearest-neighbor distance distributions largely reflected these colocalization patterns (**Figure 6F-G**). These data suggest that acetylation perturbations differentially affect how BRD4 spots associate with Mediator versus Pol II, with BRD4→POLR2A proximity surprisingly increased especially upon reduced chromatin association of BRD4. In total, these data suggest that perturbing acetylation rewires the organization of transcriptional condensates and mainly affects the BRD4 spots not associating with MED14/POLR2A spots. Specifically, HDAC inhibition leads to increase in BRD4-only spots and less association of BRD4 with Mediator or Pol II spots.

Together, these findings establish BRD4-only spots as uniquely sensitive to histone acetylation. Blocking writers or readers (less binding to chromatin) selectively disrupts BRD4-only spots—which show high chromatin enrichment but are depleted in Mediator, Pol II, and Pol II Ser5 phosphorylation (Figure 2)—whereas blocking erasers (more histone acetylation) expands this population in a bromodomain-dependent manner. These chromatin-associated, machinery-depleted condensates represent a major population under acetylation-dependent control and are distinct from previously described initiation and elongation transcriptional condensates. We propose a model (**Figure 6H**) that these BRD4-only condensates represent BRD4 clusters likely at enhancers that might be upstream of transcription initiation condensates containing Mediator and Pol II. This association of BRD4 in the latter might result from a transition into or association with initiation condensates and is seemingly less dependent on its bromodomain function.

## Discussion

By combining endogenous fluorescent tagging with high-content, time-resolved live-cell imaging and systematic chemical perturbations, we establish a quantitative framework for analyzing transcription-associated condensates under near-physiological conditions. This overcomes key limitations of prior work, in which condensate behavior has often been inferred from overexpression systems or static endpoint measurements. Such approaches have been instrumental in defining principles of phase separation and transcription hub formation but can also exaggerate condensate formation or mask compositional heterogeneity. In contrast, our platform enables direct comparison of BRD4, MED14, and POLR2A behavior at endogenous levels across thousands of perturbation conditions. The resulting dataset provides the first systematic phenotypic map of pharmacologically perturbed transcription condensates across multiple core regulators and over time.

A central finding is that BRD4, MED14, and POLR2A do not form a single uniform condensate class. Instead, they define partially overlapping populations with distinct compositions, dynamics, and drug sensitivities. This builds on earlier work showing co-condensation of BRD4, Mediator, and Pol II at super-enhancers and active genes by revealing that endogenous transcriptional assemblies are more heterogeneous than often appreciated. In particular, we identified a prominent population of BRD4-only spots that are depleted of Mediator, Pol II, and markers associated with active transcription. Importantly, their existence is itself an important finding: BRD4-rich condensates are not necessarily equivalent to canonical transcriptionally engaged Mediator/ Pol II hubs. This decoupling between BRD4 condensation and transcriptional condensate formation is consistent with studies of BRD4-driven assemblies in other contexts, including BRD4-NUT fusion oncoprotein, where condensate formation and transcriptional activation can be mechanistically separable^55^. Strom et al. demonstrated that acetylated chromatin binding thermodynamically enables BRD4 condensate nucleation below the concentration threshold required for off-chromatin condensation^56^. Our observation that BRD4-only spots are more chromatin-enriched than MED14 or POLR2A spots, and are selectively depleted by BET/HAT inhibition, provides in-cell evidence for this chromatin-seeded nucleation model operating at endogenous protein levels. Notably, ‘BRD4-only’ is an operationally defined category constrained by our detection threshold. The underlying biology may instead reflect a compositional continuum, in which condensates span a range of BRD4:MED14:Pol II stoichiometries rather than falling into discrete classes.

Our data support a model in which BRD4 acts upstream of Mediator and Pol II in condensate organization. Several observations support this interpretation. First, BRD4 spots not only exist independently of active transcription initiation machinery, their numbers also remain robust upon inhibition of transcription initiation, whereas MED14 spots are strongly depleted and BRD4-MED14 overlap decreases. Second, elongation inhibition increases BRD4 spots, indicating that BRD4 scaffold formation can be uncoupled from productive transcriptional progression. Third, BRD4-only spots are more strongly associated with chromatin than spots enriched for downstream transcriptional machinery and uniquely sensitive to the acetylation state of the chromatin. Together, these findings support a stepwise maturation model rather than a static condensate architecture: BRD4-rich spots may form first on acetylated chromatin, and a subset of these may subsequently recruit Mediator and Pol II to generate more transcriptionally engaged states. This interpretation is further supported by previous reports that showed BRD4 perturbations affecting both Mediator and Pol II clusters and aligns with a „dynamic kissing” view of transcriptional organization, in which partially overlapping mesoscale assemblies interact transiently rather than forming a constitutively mixed compartment^26^. It further is in line with observations that BRD4 constitutively occupies chromatin prior to transcriptional activation and functions as a mitotic bookmark for post-mitotic gene reactivation^57–59^.

In this setting, BRD4-only spots likely represent enhancer-centric or promoter proximal scaffold assemblies that prime loci for rapid activation. This model is supported by known domain functions of BRD4: bromodomains mediate association with acetylated chromatin, whereas intrinsically disordered regions have been implicated in multivalent interactions, condensate formation and recruitment of co-activators^29,60^. Our data suggest that bromodomain-dependent chromatin engagement is particularly important for the BRD4-only population, while transition toward Mediator- and Pol II-containing condensates may reflect a downstream maturation step. Mediator is more dependent than BRD4 on ongoing transcription initiation, which further supports the notion that BRD4 can nucleate or mark transcription-competent chromatin states upstream of full initiation hub assembly.

At the same time, we caution that our data do not yet establish whether BRD4-only spots are strictly pre-transcriptional. Alternatively, these spots persist at genes or regulatory regions between or after transcriptional bursts, thereby marking loci that are active or activatable even when not initiating at that specific moment. This possibility is compatible with recent views of transcription as a burst-like process^61^ and with reports that components of the transcriptional machinery do not need to be continuously colocalized^26,32^. It is also consistent with the observation that elongating Pol II moves away from initiation-associated spots, such that absence of POLR2A enrichment at a BRD4 spot does not necessarily imply absence of transcriptional relevance. Moreover, signal-induced release of chromatin-bound BRD4 is required for its transition to active transcriptional regulation, suggesting that chromatin-associated BRD4 can serve as a reservoir that is mobilized during transcription activation^59^. Together, these observations support a model in which at least a subset of BRD4-only spots represent chromatin-associated assemblies that exist either upstream of or in between fully engaged transcriptional states. Whether they are best described as priming condensates, inter-burst scaffold structures, or a mixture of both remains beyond the scope of this study. An alternative possibility is that some BRD4-only spots correspond to chromatin regions with predominantly non-transcriptional functions. BRD4 has been implicated in DNA double-strand break repair, chromatin organization, maintenance of higher-order chromatin structure, and response to replication stress^62–65^. Such sites could recruit BRD4 without requiring concomitant Pol II enrichment. Distinguishing between these possibilities will require future studies combining transcriptional readouts with functional perturbation and locus-specific analyses.

The compound screen identified histone acetylation as the dominant regulatory axis selectively for the BRD4-only class. BET and HAT inhibition preferentially deplete BRD4 spots, whereas HDAC inhibition expands them. This is consistent with the established role of BRD4 as a reader of acetylated histones and with evidence that disruption of BRD4 chromatin binding preferentially impairs transcription of genes associated with BRD4-enriched regulatory regions, particularly super-enhancers^12,60,66^. The effects on MED14- and POLR2A-containing spots were much weaker. Thus, histone acetylation does not merely tune overall transcriptional condensate abundance but redistributes the balance between distinct compositional states. We therefore propose a two-step model in which acetylation creates a chromatin environment that permits BD-dependent BRD4 accumulation into chromatin-enriched condensates, whereas only a subset of these subsequently transition into Mediator/Pol II-containing transcriptional condensates. In this view, acetylation gates access to a BRD4-dependent scaffold state, while additional inputs determine productive maturation.

Our time-resolved comparison of initiation and elongation inhibitors across the three factors further provides a stage-specific map of condensate dependencies. CDK7 inhibition strongly destabilizes MED14-containing assemblies while BRD4 spot numbers remained unaffected, indicating that Mediator assemblies require ongoing CTD phosphorylation and/or Pol II recruitment to stay stable. Inhibition of productive elongation increased BRD4 spots. This argues that Mediator condensates are more tightly coupled to initiation competence, while BRD4 may function as a buffering scaffold for the transcription machinery. These results support the idea that condensate composition relates to functional state: BRD4-rich, MED14-poor spots may correspond to enhancer-primed or transcription-permissive chromatin, while BRD4/MED14/POLR2A overlapping spots mark a more engaged initiation stage.

Kinase inhibitors produced weaker effects than epigenetic perturbations. One interpretation is that condensate organization is more directly governed by chromatin state than by upstream signaling pathways, at least under the tested conditions. Another possibility is that many signaling perturbations influence transcription output without strongly altering mesoscale condensate features detectable by our assay. As our readouts capture spatial organization rather than direct RNA synthesis, modest imaging phenotypes do not necessarily reflect modest transcriptional consequences. Nonetheless, the contrast between the strong acetylation-linked effects on BRD4-only condensates and weaker phenotypes for many kinase inhibitors reinforces the view that histone acetylation and BRD4 binding is a primary organizing principle of these assemblies. This does not imply that kinase signaling plays a minor role in transcription regulation. Indeed, more than 100 signal-activated kinases have been reported to phosphorylate Pol II CTD for selective regulation of Pol II function^67^. Further studies examining how signaling pathways (e.g., receptor tyrosine kinases, MAPK cascades) modulate enhancer acetylation, BRD4 and Pol II condensates will be important for understanding stimulus-responsive transcriptional regulation.

More broadly, our findings argue against viewing transcriptional condensates as a single entity. Instead, they appear to comprise a hierarchy of partially overlapping assemblies with different molecular dependencies and likely different functional roles. This is consistent with emerging models from super-resolution and live-cell studies suggesting that what appears as one diffraction-limited focus can contain multiple nanoscale substructures and transiently interacting components^26,27,32,68^. Our confocal analysis therefore likely captures mesoscale states rather than indivisible physical droplets. The observation that BRD4-only spots can be abundant, chromatin-enriched, and differentially drug-sensitive indicates that these reflect biologically meaningful states of nuclear organization. Even if some fraction reflects transient or unresolved overlap states, the selective perturbation responses indicate that they represent a distinct imaging phenotype with mechanistic relevance.

This study has several limitations. All experiments were performed in the HAP1 cancer cells, and condensate organization may differ in primary cells or across cell types, differentiation states, or physiological context. Further, endogenous tagging, while preferable to overexpression, is not entirely without caveats: the tags could affect protein behavior, and different fluorophores were used for BRD4 versus MED14/POLR2A, with different brightness and photostability. Also, conventional confocal imaging does not resolve sub-diffraction architectures, and some apparently BRD4-only spots may contain nearby or weak MED14/POLR2A signal below our detection threshold. Relatedly, co-localization assignment is necessarily imperfect and depends on segmentation and intensity thresholds. In addition, enrichment of transcription-associated markers does not establish function, and our screen does not directly measure transcriptional output, nascent RNA production across most conditions, or locus-specific gene regulation. We do not conclude that BRD4-only spots are definitively functional, pre-initiation, or non-transcriptional structures. Finally, although mechanism-of-action-based analyses help mitigate compound-specific off-target concerns, pharmacological perturbations remain indirect and should be interpreted accordingly. In addition, the compound screen was performed in a single replicate, and thus the results should be considered exploratory and require independent validation, as demonstrated for the HAT and HDAC inhibitors.

Despite these limitations, the conceptual implications are substantial. The identification of a large endogenous BRD4-only condensate population reveals an unappreciated layer of heterogeneity in nuclear transcriptional organization. Whether these spots represent enhancer-priming structures, inter-burst bookmarks, or another chromatin-associated BRD4 state, they point to a model in which BRD4 condensation is not synonymous with transcription itself but instead defines a separable regulatory layer upstream of or adjacent to canonical Mediator/Pol condensates. This suggests that nuclear condensate organization can encode regulatory potential, not just ongoing activity. Such a model has also therapeutic implications: targeting BD-dependent BRD4 assemblies may selectively reshape enhancer-centric chromatin condensates without uniformly dismantling all transcription machinery, an idea that may especially be relevant in diseases driven by aberrant super-enhancer activity, as has been reported in cancer^10,66,69,70^.

In summary, we establish a scalable live-cell phenomics platform for condensate biology that is broadly applicable across condensate types and reveal that endogenous transcriptional condensates are compositionally heterogeneous and dynamically regulated. BRD4-only condensates define a distinct chromatin-enriched class depleted of Mediator and RNA polymerase II. Chemical perturbation screening identifies histone acetylation as a dominant regulatory axis, with BET and HAT inhibition selectively depleting BRD4-only condensates and HDAC inhibition expanding them. We propose that these assemblies represent a BRD4-dependent chromatin scaffold state that can persist independently of transcription initiation and, in some contexts, mature into Mediator- and Pol II-containing transcription hubs. More broadly, our findings establish condensate composition as a quantitative and mechanistically informative phenotype that encodes regulatory state within the nucleus.

## Materials and Methods

### Cell lines and culture

Diploid HAP1 cells, a human cell line derived from the chronic myelogenous leukemia line KBM-7, were maintained at 37°C in a humidified incubator with 5% CO₂. Cells were cultured in Iscove’s Modified Dulbecco’s Medium (IMDM; Gibco, Waltham, USA) supplemented with 10% (v/v) fetal bovine serum (Gibco, FBS, qualified, Brazil lot #2307587 or Value FBS lot #2529121) and passaged approximately twice per week at 70–80% confluence using 0.25% trypsin-EDTA (Gibco).

### Endogenous tagging

Endogenous tagging was performed using the CRISPaint (CRISPR-assisted insertion tagging) method. Starting from a HAP1 BRD4::eGFP cell line (gift from Stefan Kubicek, Vienna, Austria), the genes *MED14, POLR2A,* and *SMARCA4* were sequentially tagged at their endogenous C-termini. Target-specific guide RNAs were cloned into pSPgRNA plasmids (Addgene #47108) and co-transfected with frame-selector and donor plasmids at a 1:1:2 mass ratio using X-tremeGENE 9 (Sigma-Aldrich, Darmstadt, Germany). Donor plasmids encoded TagRFP-T2A-PuroR (Addgene #80971) or SNAP-dTAG-HA-T2A-NeoR (generated in-house), with antibiotic resistance cassettes separated by T2A self-cleaving peptides. Following 48 hours of transfection, cells underwent selection with 1 µg/ml puromycin (InvivoGen, San Diego, USA) or 1000 µg/ml G418/neomycin (Sigma-Aldrich). Surviving clones were validated by genomic PCR, Western blotting, and fluorescence microscopy.

### Live-cell imaging

Automated high-content confocal imaging was performed on an Opera Phenix High Content Screening System (PerkinElmer, Waltham, USA) equipped with spinning-disk confocal optics. Cells were seeded in poly-D-lysine-coated (Merck, Darmstadt, Germany) 96-well or 384-well PhenoPlates (Revvity, Waltham, USA) at 20,000 or 5,000 cells/well, respectively, 24 hours prior to imaging. Before imaging, cells were labeled with 1 µg/ml Hoechst 33342 (Thermo Fisher Scientific, Waltham, USA) for 30 minutes at 37°C, washed three times, and incubated for an additional 30 minutes. For SMARCA4 visualization, cells were additionally labeled with 250 nM SNAP-Cell 647-SiR (New England Biolabs, Ipswich, USA).

Imaging was performed at 37°C with 5% CO₂ using a 63× water-immersion objective (1.15 NA) with 2×2 binning (188 nm pixel size). Three z-planes were acquired with 2 µm spacing. Time-lapse imaging was conducted over 0–12 hours at multiple timepoints (0, 1, 2, 3, 4, 8, and 12 hours). Channel-specific settings were maintained constant: Hoechst 33342 (405 nm excitation, 435–480 nm emission, 120 ms, 60% power), eGFP (488 nm, 500–550 nm, 700 ms, 70% power), TagRFP (561 nm, 570–630 nm, 700 ms, 70% power), and Alexa Fluor 647 (640 nm, 650–760 nm, 500 ms, 50% power).

### Image analysis

Automated image analysis was performed using Harmony software v4.9 (PerkinElmer). Maximum intensity projections were generated from z-stacks following flatfield correction. Nuclei were segmented in the Hoechst 33342 channel using standard morphological filters (area >75 µm², roundness >0.75), and border objects were excluded. Spot detection was performed using Harmony software’s ‘Find Spots’ algorithm (Method A) within Hoechst-defined nuclei. Marker-specific intensity thresholds were: BRD4::eGFP 0.04, MED14::TagRFP 0.05, POLR2A::TagRFP 0.06. Splitting sensitivity was set to 1 (no extra splitting) for all channels. Extracted features included spot count, diameter, intensity, colocalization (based on center-to-center Euclidean distances), nearest-neighbor distances, and marker enrichment (mean spot intensity divided by mean nuclear intensity). Single-cell and single-spot quantification data were exported for downstream analysis.

### Immunofluorescence

Following live-cell time courses, cells were fixed with 4% paraformaldehyde (Sigma-Aldrich) in DPBS for 15 minutes at room temperature, quenched with 50 mM Tris/100 mM NaCl, permeabilized with 0.2% Triton X-100 for 20 minutes, and blocked with 3% BSA. Primary antibodies targeting RNA polymerase II CTD (total: Abcam ab26721, 1:1000; Ser5-P: Abcam ab5408, 1:1000; Ser2-P: Sigma-Aldrich 04-1571, 1:1000) were incubated for 1 hour at room temperature. Samples were washed and incubated with Alexa Fluor 647-conjugated secondary antibodies (Invitrogen, 1:500) for 1 hour, then imaged using identical acquisition settings as pre-fixation imaging. For nascent RNA detection, cells were pulse-labeled with 1 mM 5-ethynyl uridine (5-EU; Sigma-Aldrich) for 60 minutes, fixed, and subjected to click-chemistry with 5 µM Alexa Fluor 647 azide using the Click-iT RNA Imaging Kit (Invitrogen, C10330).

### FRAP experiments

Fluorescence recovery after photobleaching (FRAP) was performed on a VisiScope spinning-disk confocal microscope (Visitron Systems, Puchheim, Germany) with a 63× oil-immersion objective and 2×2 binning. HAP1 BRD4::eGFP cells were seeded in 8-well µ-Slide chamber slides (Ibidi) and imaged with 488 nm excitation (300 ms, 70% power). Individual BRD4 spots (three-pixel radius) were photobleached using the 488 nm laser at 20% power with five bursts (100 ms/pixel dwell time). Recovery was tracked over 60 frames at 1-second intervals following 10 pre-bleach frames. Data were analyzed using the easyFRAP software^71^ with full-scale normalization and single-exponential curve fitting to extract mobile fractions and recovery half-times (t½).

### Super-resolution microscopy

Single-molecule localization microscopy (SMLM) was performed on a NanoImager microscope (ONI, Oxford, UK). Fixed and permeabilized cells were immunostained with anti-GFP conjugated to Alexa Fluor 647 (NanoTag Biotechnologies N0301-AF647-L, 1:500) and imaged in GLOX/BME switching buffer (50 mM Tris-HCl pH 8, 10 mM NaCl, 500 µg/ml glucose oxidase, 40 µg/ml catalase, 10% glucose, 50 mM β-mercaptoethanol). A total of 20,000 frames were acquired under continuous 647 nm excitation. Localization data were processed using the ThunderSTORM plugin^72^ in Fiji/ImageJ^73^ with B-spline wavelet filtering, local-maximum detection, and integrated-Gaussian PSF fitting with maximum likelihood estimation. Localizations were drift-corrected, filtered for xy-uncertainty (80–300 nm), and rendered at 5× magnification.

### Compound perturbations

Compounds were dissolved in DMSO at 10 mM stock concentrations and stored at −80°C. Working concentrations were: THZ1 (CDK7 inhibitor, Hycultec, HY-80013; 100 nM), Flavopiridol (CDK9 inhibitor, Hycultec, HY-10006; 100 nM), JQ1 (BET inhibitor, Sigma-Aldrich, SML0974; 500 nM and 1 µM), A485 (p300/CBP HAT inhibitor, Hycultec, HY-107455; 100 nM and 500 nM), and SAHA/vorinostat (HDAC inhibitor, Sigma-Aldrich, SML0061; 100 nM and 500 nM). DMSO vehicle controls were included at equivalent volumes (≤0.01% v/v). Following cell labeling, medium was replaced with compound-containing medium, and cells were immediately imaged.

### High-throughput compound screen

A library of >1,000 compounds spanning epigenetic modulators, kinase inhibitors, anti-cancer agents, and CLOUD collections was screened in 384-well format. Compounds were pre-arrayed and resuspended in phenol red-free IMDM immediately before treatment. Liquid handling for compound dispensing and medium exchange was performed using an Integra VIAFLO 384 electronic pipetting system (Integra Biosciences, Hudson, USA). Each compound was tested at two concentrations with plate-matched DMSO controls and positive controls on every plate. Quality control filters excluded wells with low nuclei counts (<10 nuclei/well), cytotoxicity (>80% reduction in nuclei at 12 hours), autofluorescence (top and bottom 1% quantiles), or intensity outliers (top and bottom 1% quantiles). Across retained plates, 8.6% of wells (247/2,900) failed at least one QC criterion and were excluded.

### MOA annotation

To standardize mechanism of action (MOA) annotations across diverse compound libraries, we implemented a multi-step computational workflow. Compound identities were first canonicalized by resolving compound names and/or SMILES (Simplified Molecular Input Line Entry System) chemical structure strings via PubChem and cross-referenced to unique ChEMBL database identifiers. For each compound, molecular target annotations were extracted from curated ChEMBL mechanism entries when available. For compounds lacking curated annotations, targets were inferred from ChEMBL bioactivity data based on defined activity thresholds.

### Data analysis compound screen

Feature values were robustly z-score normalized relative to plate-matched DMSO distributions: z = (x − median_DMSO) / (1.4826 × MAD_DMSO), where MAD is the median absolute deviation. Time-course trajectories were summarized by calculating the signed area under the curve (AUC), integrating z-scores over time. Feature correlations were computed as pairwise Pearson coefficients. Principal component analysis (PCA) was performed on mean AUC values using column-centered data. Hierarchical clustering used Pearson correlation distances (1 − r) with average linkage. Hit calling identified compounds in the top or bottom 1% by feature. MOA enrichment was calculated as the fraction of a given MOA class among hits divided by its fraction among all screened conditions. Statistical analyses and visualization were performed in R v4.5.1 using RStudio^74,75^ and ggplot2^76^.

## Data and Code availability

The codes for the analyses are available at https://github.com/SchickLab/Transcriptional-condensate-characterization-and-screen/.

## Author contributions

S.S. conceived and supervised the study, S.S. and S.Sh. designed and coordinated the study. S.Sh., A.S., A.-S.S. performed experiments and analyses, K.H. performed data analyses, S.Sh. performed computational analyses, M.G. supported microscopy experiments, and F.K. supported statistical analyses. S.S. and S.Sh. wrote the manuscript. All authors reviewed and approved the manuscript. S.Sh., T.S. and S.S. acquired funding.

## Ethics / conflicts of interest

The authors declare no competing interests.

## Acknowledgements

We thank all members of the Schick Lab for useful discussions throughout the study. We thank Stefan Kubicek (CeMM, Vienna, Austria) for providing the HAP1 BRD4::eGFP cell line and Stefan Kubicek and Anna Koren (CeMM, Vienna, Austria) for assistance with the compound screen library. We thank Sandra Ritz, Petri Turunen, and Anusha Koplan of the IMB Microscopy Core Facility for their support and assistance with high-throughput live-cell confocal imaging on the Opera Phenix (DFG project #316215830) and with live-cell time-lapse imaging and FRAP on the Visitron Spinning Disc (DFG project #402386039). We thank the IMB Media Lab for technical support. We also thank Sina Wittmann (IMB, Mainz) for her helpful input and discussion during manuscript preparation. Portions of the manuscript were edited for clarity using AI-assisted language tools. This project was funded by the Deutsche Forschungsgemeinschaft (DFG, German Research Foundation) - SFB 1551 – Project No. 464588647 and supported by the Max Planck Graduate Center with the Johannes Gutenberg-Universität Mainz (MPGC). The Schick laboratory is additionally supported by the Deutsche Forschungsgemeinschaft (DFG, German Research Foundation) – Project-ID 393547839 – SFB 1361, an ERC Starting Grant (SWItchFate, Grant 101164641), and another project was funded during the research time by the Fritz Thyssen Foundation. We thank all members of the Schick laboratory for helpful discussions.

---

**Supplementary Figure 1:**
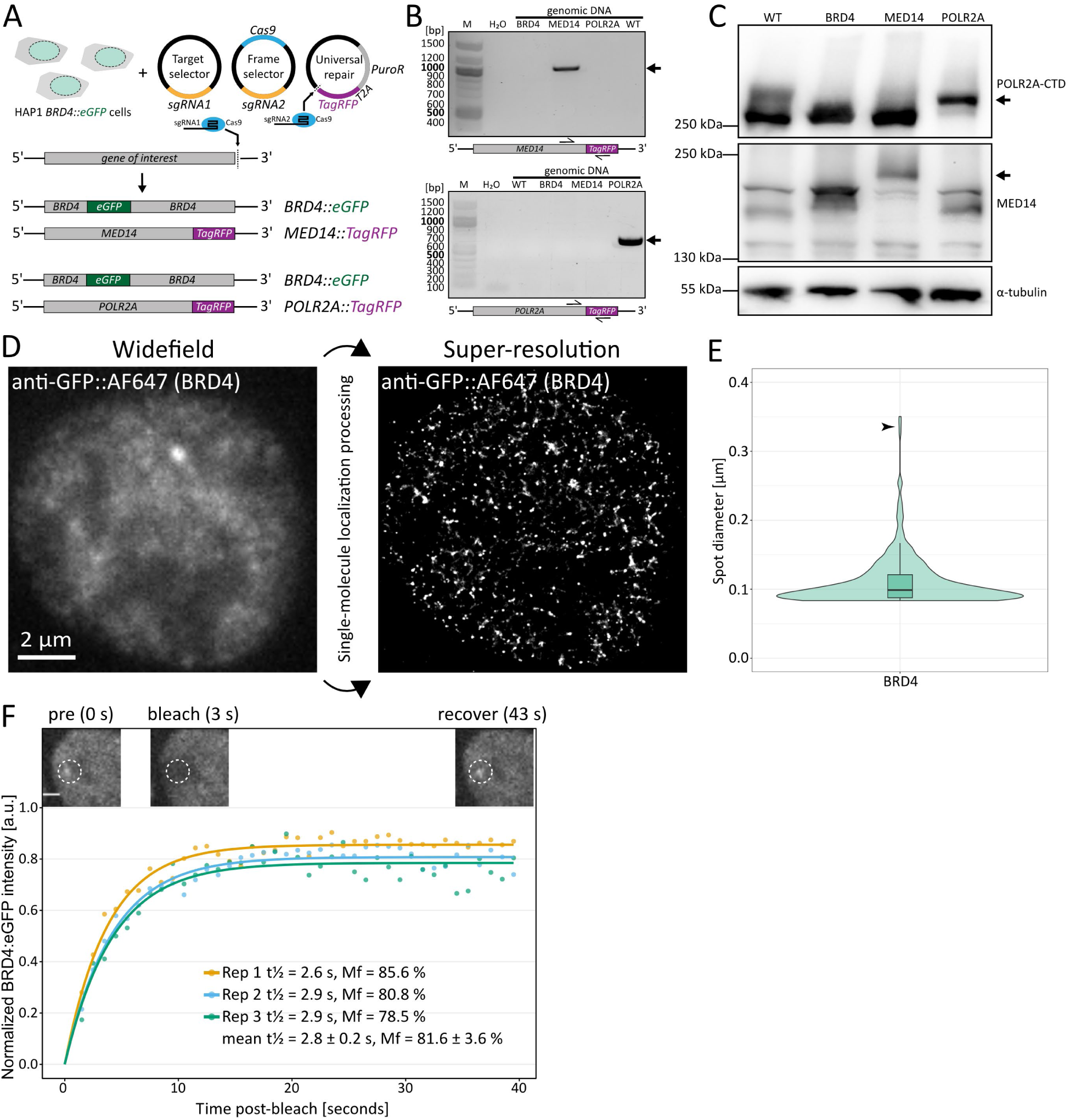
Generation and validation of endogenously tagged BRD4, MED14, and POLR2A reporter cell lines and spot spot properties. **(A)** Schematic of CRISPR-assisted insertion tagging strategy used to generate double-knockin HAP1 reporter lines. Starting from a parental BRD4::eGFP line, sgRNAs targeting the C-terminus of MED14 or POLR2A were co-delivered with donor cassettes encoding TagRFP and a puromycin resistance marker to produce BRD4::eGFP; MED14::TagRFP and BRD4::eGFP; POLR2A::TagRFP lines (see Methods). **(B)** Genotyping PCR of clones used for downstream experiments. Top: PCR spanning MED14 edit shows product of expected size (arrow). Bottom: PCR spanning POLR2A edits shows a product of expected size (arrow). M, marker; H2O, no-template control; WT, unedited parental control. Water-only, no template (H2O), Wildtype (WT), BRD4::eGFP (BRD4), BRD4::eGFP_MED14::TagRFP (MED14), BRD4::eGFP_MED14::TagRFP (POLR2A) samples indicated. **(C)** Western blot validation of tagged lines. Top: anti-POLR2A (CTD) shows expected shift in POLR2A::TagRFP lane (arrow). Middle: anti-MED14 detects MED14::TagRFP band (arrow). Bottom: α-tubulin loading control. Representative blots from independent biological replicates are shown. Cell line samples as in (B). **(D)** Representative single nucleus images of BRD4 immunofluorescence comparing widefield (left) and single-molecule localization super-resolution reconstruction (right) of anti-GFP::AF647 signal in BRD4::eGFP cells; scale bar, 2 µm. Single-molecule processing reveals numerous sub-diffraction localizations within confocal spots. **(E)** Distribution of BRD4 spot diameters measured from super-resolution localizations (violin plot with box showing median and interquartile range). Median spot diameter is ∼0.10 µm (arrowhead indicates median). **(F)** Fluorescence recovery after photobleaching (FRAP) of BRD4::eGFP foci in live cells treated with 0.01% DMSO (vehicle control). Individual foci were bleached and fluorescence recovery was monitored over time. Mean fluorescence intensity was background-subtracted, corrected for acquisition bleaching using an unbleached reference ROI, and normalized to the pre-bleach intensity (= 1.0). Data points represent the mean normalized fluorescence per biological replicate (FRAP4, n = 9; FRAP5, n = 11; FRAP7, n = 14 foci). Solid lines show single exponential fits. Recovery half-time (t½ = ln(2)/k) and mobile fraction (Mf) were extracted per replicate and are then again reported as mean ± SD across three experiments.

**Supplementary Figure S2:**
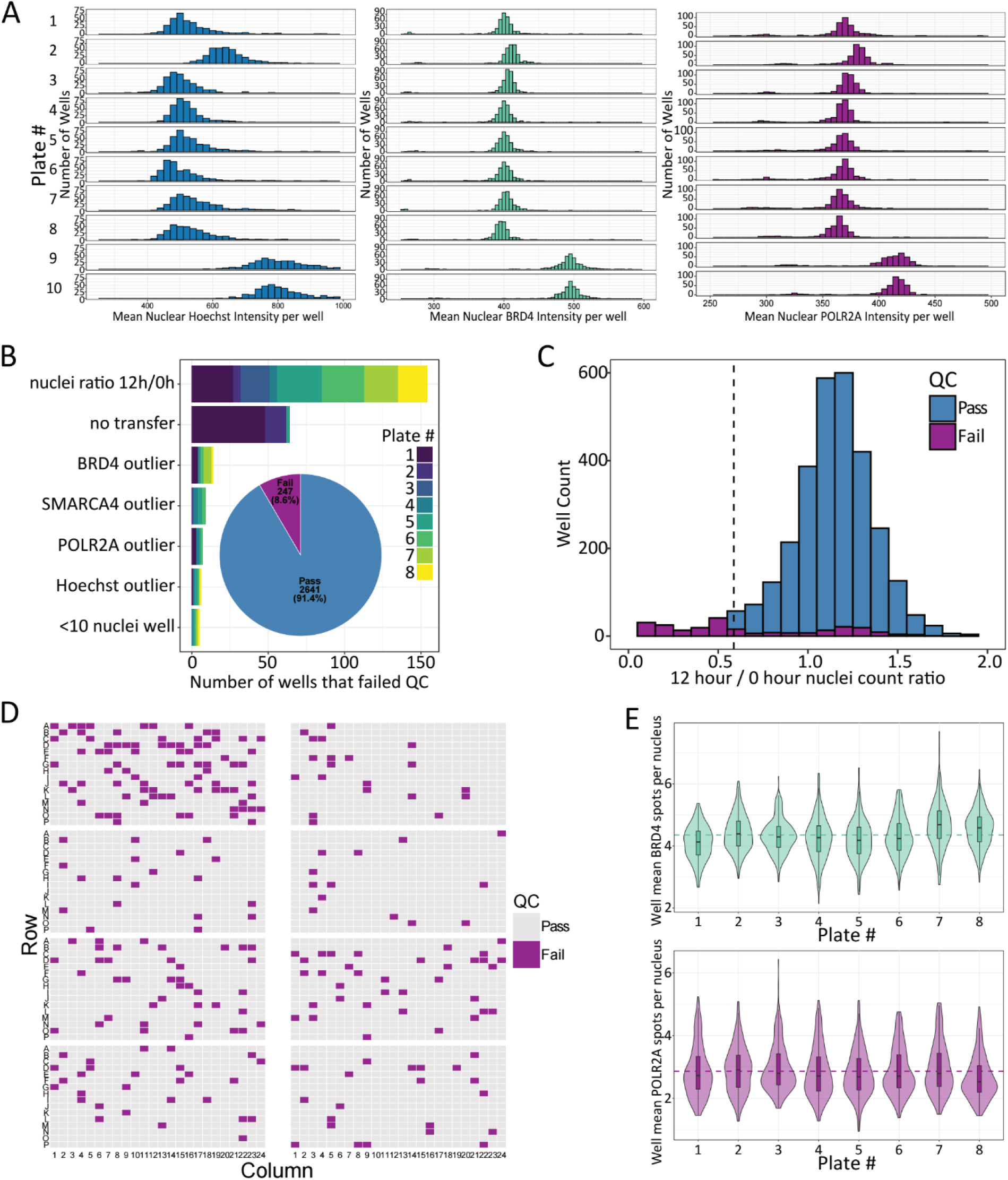
Quality control (QC) of the high-content imaging screen across plates and wells. **(A)** Plate-wise distributions of mean nuclear Hoechst intensity, BRD4 intensity, and POLR2A intensity per well across the ten screened plates. Each row corresponds to a plate, and each histogram summarizes the well-level distribution for the indicated nuclear feature. **(B)** Summary of quality control failures across the full compound screen. The stacked bar plot shows the number of wells failing each QC criterion, grouped by plate, and the pie chart indicates the overall fraction of wells passing versus failing QC. Wells were excluded for low nuclei ratio at 12 h versus 0 h, no compound transfer, BRD4 outliers, SMARCA4 outliers, POLR2A outliers, Hoechst outliers, or fewer than 10 nuclei per well. **(C)** Distribution of 12 h/0 h nuclei count ratio across all wells, separated into pass and fail categories. The dashed line marks the QC threshold used to identify toxic or low-coverage wells. **(D)** Plate map of well-level QC outcomes across the screened 384-well plates. Wells failing at least one QC criterion are shown in purple, and passing wells are shown in gray. The layout shows a uniform spatial distribution of failures without strong well-position bias. **(E)** Per-plate violin plots of mean BRD4 spot count per nucleus and mean POLR2A spot count per nucleus for wells passing QC. Dashed horizontal lines indicate the DMSO baseline median for each plate, showing comparable plate-to-plate spot distributions after QC filtering.

**Supplementary Figure S3:**
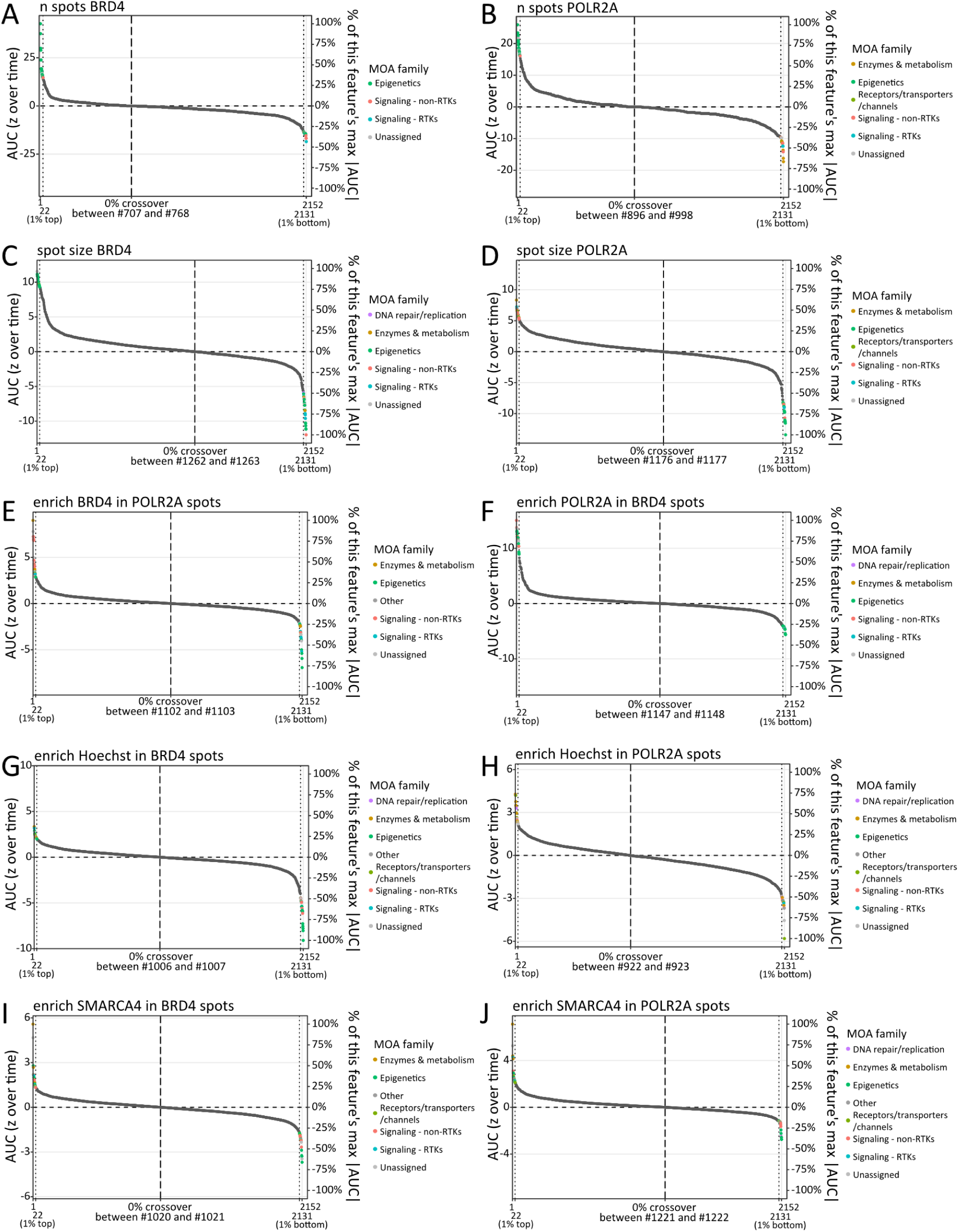
Ranked response distributions of all ten imaging features across the compound screen. (A–J) AUC values for each feature plotted against compound rank. Positive AUC values represent increases and negative values decreases relative to DMSO. Dashed vertical lines indicate 1 % thresholds used for hit calling. Right-hand y-axis shows cumulative % of hits.

**Supplementary Figure 4:**
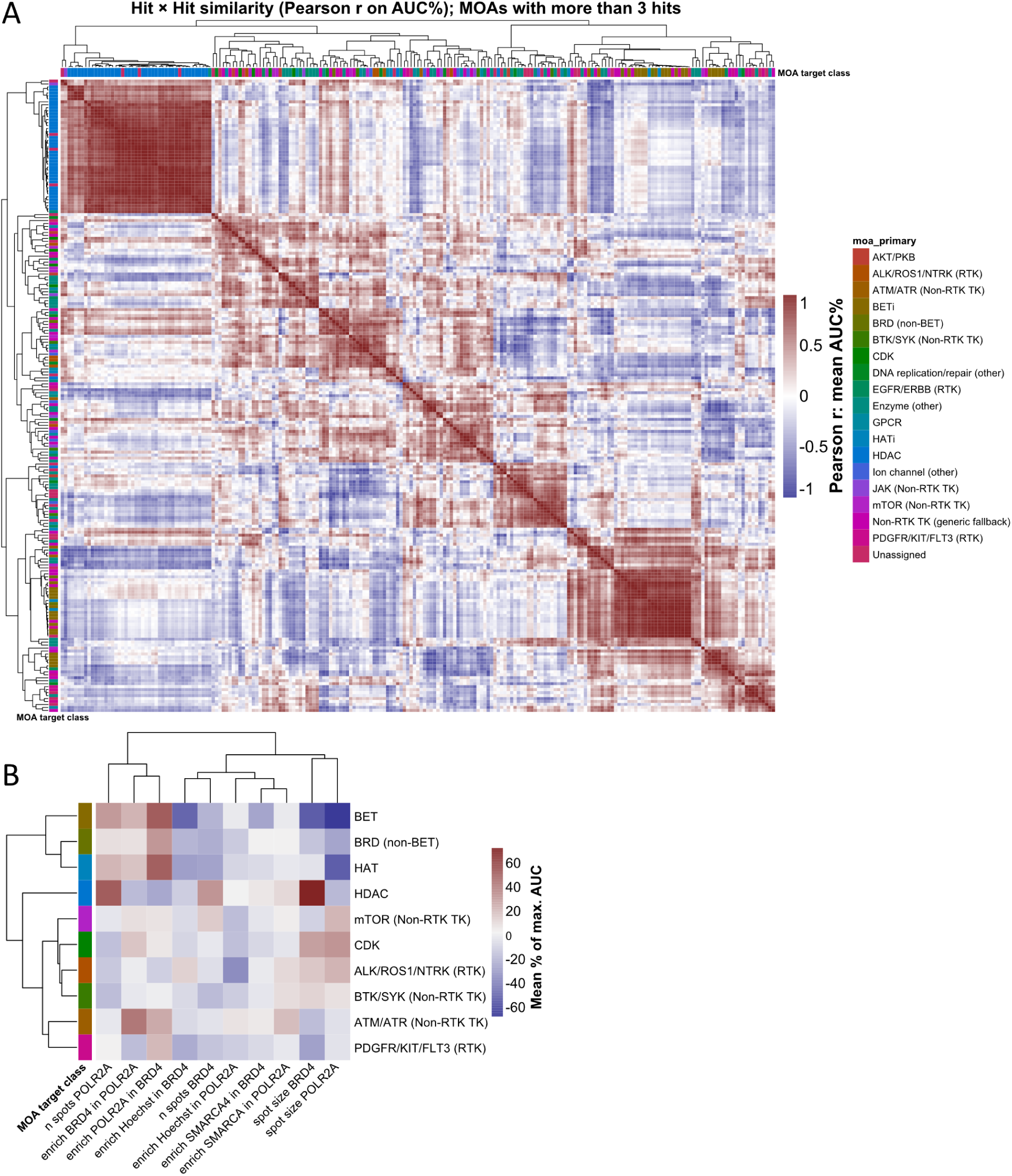
Compound-level hit similarity and MOA feature signatures recapitulate mechanism-of-action-based condensate phenotype clusters. (A) Hit × Hit similarity matrix showing pairwise Pearson correlations of AUC% profiles across all ten imaging features for individual compound-concentration conditions classified as hits (top or bottom 1% in at least one feature; n = 242 hit conditions). Rows and columns are ordered by unsupervised hierarchical clustering (Pearson correlation distance, average linkage). Color bar indicates MOA target class annotation for each hit condition. Color scale: Pearson r on mean AUC%, ranging from −1 (blue) to +1 (red). (B) Heatmap of mean AUC% (% of maximum AUC) for each MOA target class (rows) across the ten condensate imaging features (columns), restricted to MOA classes with more than three hit conditions. Rows and columns are ordered by unsupervised hierarchical clustering. Color scale ranges from −60 to +60% of maximum AUC. Features shown are spot counts (n spots) and spot sizes for BRD4 and POLR2A, co-enrichment of BRD4 within POLR2A spots and vice versa, Hoechst chromatin enrichment within BRD4 and POLR2A spots, and SMARCA4 enrichment within BRD4 and POLR2A spots.

## References

1. Guo, J. Transcription: the epicenter of gene expression. J. Zhejiang Univ. Sci. B 15, 409–411 (2014).

2. Sperling, S. Transcriptional regulation at a glance. BMC Bioinformatics 8, S2 (2007).

3. Jovanovic, M. et al. Dynamic profiling of the protein life cycle in response to pathogens. Science 347, 1259038 (2015).

4. Marguerat, S., Lawler, K., Brazma, A. & Bähler, J. Contributions of transcription and mRNA decay to gene expression dynamics of fission yeast in response to oxidative stress. RNA Biology 11, 702–714 (2014).

5. Rabani, M. et al. Metabolic labeling of RNA uncovers principles of RNA production and degradation dynamics in mammalian cells. Nat Biotechnol 29, 436–442 (2011).

6. Ptashne, M. & Gann, A. Genes & Signals. (Cold Spring Harbor Laboratory Press, Cold Spring Harbor, New York, 2002).

7. Reiter, F., Wienerroither, S. & Stark, A. Combinatorial function of transcription factors and cofactors. Curr Opin Genet Dev 43, 73–81 (2017).

8. Spitz, F. & Furlong, E. E. M. Transcription factors: from enhancer binding to developmental control. Nat Rev Genet 13, 613–626 (2012).

9. Parker, S. C. J. et al. Chromatin stretch enhancer states drive cell-specific gene regulation and harbor human disease risk variants. Proc. Natl. Acad. Sci. U.S.A. 110, 17921–17926 (2013).

10. Hnisz, D. et al. Super-Enhancers in the Control of Cell Identity and Disease. Cell 155, 934–947 (2013).

11. Whyte, W. A. et al. Master Transcription Factors and Mediator Establish Super-Enhancers at Key Cell Identity Genes. Cell 153, 307–319 (2013).

12. Lovén, J. et al. Selective Inhibition of Tumor Oncogenes by Disruption of Super-Enhancers. Cell 153, 320–334 (2013).

13. Wang, Z. et al. Genome-wide Mapping of HATs and HDACs Reveals Distinct Functions in Active and Inactive Genes. Cell 138, 1019–1031 (2009).

14. Creyghton, M. P. et al. Histone H3K27ac separates active from poised enhancers and predicts developmental state. Proc. Natl. Acad. Sci. U.S.A. 107, 21931–21936 (2010).

15. Qian, H. et al. Super-enhancers and the super-enhancer reader BRD4: tumorigenic factors and therapeutic targets. Cell Death Discov. 9, 470 (2023).

16. Devaiah, B. N. et al. BRD4 is a histone acetyltransferase that evicts nucleosomes from chromatin. Nat Struct Mol Biol 23, 540–548 (2016).

17. Zheng, B. et al. Distinct layers of BRD4-PTEFb reveal bromodomain-independent function in transcriptional regulation. Molecular Cell 83, 2896–2910.e4 (2023).

18. Altendorfer, E., Mochalova, Y. & Mayer, A. BRD4: a general regulator of transcription elongation. Transcription 13, 70–81 (2022).

19. Jackson, D. A., Hassan, A. B., Errington, R. J. & Cook, P. R. Visualization of focal sites of transcription within human nuclei. The EMBO Journal 12, 1059–1065 (1993).

20. Cisse, I. I. et al. Real-Time Dynamics of RNA Polymerase II Clustering in Live Human Cells. Science 341, 664–667 (2013).

21. Lambert, S. A. et al. The Human Transcription Factors. Cell 172, 650–665 (2018).

22. Sigler, P. B. Acid blobs and negative noodles. Nature 333, 210–212 (1988).

23. Fuxreiter, M. et al. Malleable machines take shape in eukaryotic transcriptional regulation. Nat Chem Biol 4, 728–737 (2008).

24. Lyons, H. et al. Functional partitioning of transcriptional regulators by patterned charge blocks. Cell 186, 327–345.e28 (2023).

25. Hnisz, D., Shrinivas, K., Young, R. A., Chakraborty, A. K. & Sharp, P. A. A Phase Separation Model for Transcriptional Control. Cell 169, 13–23 (2017).

26. Cho, W.-K. et al. Mediator and RNA polymerase II clusters associate in transcription-dependent condensates. Science 361, 412–415 (2018).

27. Chong, S. et al. Imaging dynamic and selective low-complexity domain interactions that control gene transcription. Science 361, eaar2555 (2018).

28. Boija, A. et al. Transcription Factors Activate Genes through the Phase-Separation Capacity of Their Activation Domains. Cell 175, 1842–1855.e16 (2018).

29. Sabari, B. R. et al. Coactivator condensation at super-enhancers links phase separation and gene control. Science 361, eaar3958 (2018).

30. Schmid-Burgk, J. L., Höning, K., Ebert, T. S. & Hornung, V. CRISPaint allows modular base-specific gene tagging using a ligase-4-dependent mechanism. Nat Commun 7, 12338 (2016).

31. Jaeger, M. G. et al. Selective Mediator dependence of cell-type-specifying transcription. Nat Genet 52, 719–727 (2020).

32. Guo, Y. E. et al. Pol II phosphorylation regulates a switch between transcriptional and splicing condensates. Nature 572, 543–548 (2019).

33. Lu, H. et al. Phase-separation mechanism for C-terminal hyperphosphorylation of RNA polymerase II. Nature 558, 318–323 (2018).

34. Pancholi, A. et al. RNA polymerase II clusters form in line with surface condensation on regulatory chromatin. Molecular Systems Biology 17, e10272 (2021).

35. Corden, J. L. Tails of RNA polymerase II. Trends in Biochemical Sciences 15, 383–387 (1990).

36. Heidemann, M., Hintermair, C., Voß, K. & Eick, D. Dynamic phosphorylation patterns of RNA polymerase II CTD during transcription. Biochimica et Biophysica Acta (BBA) - Gene Regulatory Mechanisms 1829, 55–62 (2013).

37. Harlen, K. M. & Churchman, L. S. The code and beyond: transcription regulation by the RNA polymerase II carboxy-terminal domain. Nat Rev Mol Cell Biol 18, 263–273 (2017).

38. Pancholi, A. et al. RNA polymerase II clusters form in line with surface condensation on regulatory chromatin. Mol Syst Biol 17, MSB202110272 (2021).

39. Kwiatkowski, N. et al. Targeting transcription regulation in cancer with a covalent CDK7 inhibitor. Nature 511, 616–620 (2014).

40. Chipumuro, E. et al. CDK7 Inhibition Suppresses Super-Enhancer-Linked Oncogenic Transcription in MYCN-Driven Cancer. Cell 159, 1126–1139 (2014).

41. Chao, S.-H. & Price, D. H. Flavopiridol Inactivates P-TEFb and Blocks Most RNA Polymerase II Transcription in Vivo. Journal of Biological Chemistry 276, 31793–31799 (2001).

42. Chen, R., Keating, M. J., Gandhi, V. & Plunkett, W. Transcription inhibition by flavopiridol: mechanism of chronic lymphocytic leukemia cell death. Blood 106, 2513–2519 (2005).

43. Licciardello, M. P. et al. A combinatorial screen of the CLOUD uncovers a synergy targeting the androgen receptor. Nat Chem Biol 13, 771–778 (2017).

44. Schick, S. et al. Systematic characterization of BAF mutations provides insights into intracomplex synthetic lethalities in human cancers. Nat Genet 51, 1399–1410 (2019).

45. Schick, S. et al. Acute BAF perturbation causes immediate changes in chromatin accessibility. Nat Genet 53, 269–278 (2021).

46. Varga, J., Kube, M., Luck, K. & Schick, S. The BAF chromatin remodeling complexes: structure, function, and synthetic lethalities. Biochemical Society Transactions 49, 1489–1503 (2021).

47. Clapier, C. R., Iwasa, J., Cairns, B. R. & Peterson, C. L. Mechanisms of action and regulation of ATP-dependent chromatin-remodelling complexes. Nat Rev Mol Cell Biol 18, 407–422 (2017).

48. Patil, A. et al. A disordered region controls cBAF activity via condensation and partner recruitment. Cell 186, 4936–4955.e26 (2023).

49. Davis, R. B., Kaur, T., Moosa, M. M. & Banerjee, P. R. FUS oncofusion protein condensates recruit mSWI/SNF chromatin remodeler via heterotypic interactions between prion-like domains. Protein Science 30, 1454–1466 (2021).

50. Zolotarev, N. et al. Regularly spaced tyrosines in EBF1 mediate BRG1 recruitment and formation of nuclear subdiffractive clusters. Genes Dev. 38, 4–10 (2024).

51. Mendez, D. et al. ChEMBL: towards direct deposition of bioassay data. Nucleic Acids Research 47, D930–D940 (2019).

52. Filippakopoulos, P. et al. Selective inhibition of BET bromodomains. Nature 468, 1067–1073 (2010).

53. Lasko, L. M. et al. Discovery of a selective catalytic p300/CBP inhibitor that targets lineage-specific tumours. Nature 550, 128–132 (2017).

54. Richon, V. M. et al. A class of hybrid polar inducers of transformed cell differentiation inhibits histone deacetylases. Proc. Natl. Acad. Sci. U.S.A. 95, 3003–3007 (1998).

55. Kosno, M., Currie, S. L., Kumar, A., Xing, C. & Rosen, M. K. Molecular features driving condensate formation and gene expression by the BRD4-NUT fusion oncoprotein are overlapping but distinct. Sci Rep 13, 11907 (2023).

56. Strom, A. R., et al. Interplay of condensation and chromatin binding underlies BRD4 targeting. MBoC 35, ar88 (2024).

57. Dey, A., Nishiyama, A., Karpova, T., McNally, J. & Ozato, K. Brd4 Marks Select Genes on Mitotic Chromatin and Directs Postmitotic Transcription. MBoC 20, 4899–4909 (2009).

58. Strom, A. R., et al. Interplay of condensation and chromatin binding underlies BRD4 targeting. MBoC 35, ar88 (2024).

59. Ai, N. et al. Signal-induced Brd4 release from chromatin is essential for its role transition from chromatin targeting to transcriptional regulation. Nucleic Acids Research 39, 9592–9604 (2011).

60. Filippakopoulos, P. et al. Selective inhibition of BET bromodomains. Nature 468, 1067–1073 (2010).

61. Rodriguez, J. & Larson, D. R. Transcription in Living Cells: Molecular Mechanisms of Bursting. Annual Review of Biochemistry 89, 189–212 (2020).

62. Zhang, J. et al. BRD4 facilitates replication stress-induced DNA damage response. Oncogene 37, 3763–3777 (2018).

63. Wang, R., Li, Q., Helfer, C. M., Jiao, J. & You, J. Bromodomain Protein Brd4 Associated with Acetylated Chromatin Is Important for Maintenance of Higher-order Chromatin Structure. Journal of Biological Chemistry 287, 10738–10752 (2012).

64. Floyd, S. R. et al. The bromodomain protein Brd4 insulates chromatin from DNA damage signalling. Nature 498, 246–250 (2013).

65. Barrows, J. K. et al. BRD4 promotes resection and homology-directed repair of DNA double-strand breaks. Nat Commun 13, 3016 (2022).

66. Chapuy, B. et al. Discovery and Characterization of Super-Enhancer-Associated Dependencies in Diffuse Large B Cell Lymphoma. Cancer Cell 24, 777–790 (2013).

67. Dabas, P. et al. Direct targeting and regulation of RNA polymerase II by cell signaling kinases. Science 390, eads7152 (2025).

68. Mir, M. et al. Dynamic multifactor hubs interact transiently with sites of active transcription in Drosophila embryos. Elife 7, e40497 (2018).

69. Lovén, J. et al. Selective Inhibition of Tumor Oncogenes by Disruption of Super-Enhancers. Cell 153, 320–334 (2013).

70. Bradner, J. E., Hnisz, D. & Young, R. A. Transcriptional Addiction in Cancer. Cell 168, 629–643 (2017).

71. Rapsomaniki, M. A. et al. easyFRAP: an interactive, easy-to-use tool for qualitative and quantitative analysis of FRAP data. Bioinformatics 28, 1800–1801 (2012).

72. Ovesný, M., Křížek, P., Borkovec, J., Švindrych, Z. & Hagen, G. M. ThunderSTORM: a comprehensive ImageJ plug-in for PALM and STORM data analysis and super-resolution imaging. Bioinformatics 30, 2389–2390 (2014).

73. Schindelin, J., et al. Fiji: an open-source platform for biological-image analysis. Nat Methods 9, 676–682 (2012).

74. R Core Team. R: A Language and Environment for Statistical Computing. R Foundation for Statistical Computing (2025).

75. Posit team. RStudio: Integrated Development Environment for R. Posit Software, PBC (2025).

76. Hadley Wickham. ggplot2: Elegant Graphics for Data Analysis. (2016).

